# Comparative ASCL1 interactome analysis reveals CDK2-Cyclin A2 as suppressors of differentiation in MYCN-amplified neuroblastoma

**DOI:** 10.1101/2025.11.11.687869

**Authors:** Lidiya Mykhaylechko, Laura M. Woods, Evangelia K. Papachristou, Roshna L. Gomez, Revathy Ramachandran, Jethro Lundie-Brown, Daniel Marcos, Rosalind Drummond, Fahad R. Ali, Jason Carroll, Anna Philpott

## Abstract

Neuroblastoma is a heterogeneous paediatric cancer arising from developmentally arrested neuronal precursors, where restoring differentiation offers therapeutic promise. ASCL1, a pro-neural transcription factor, is widely expressed in neuroblastoma and can drive either proliferation or differentiation depending on the cellular context. Here, we show that distinct MYCN-amplified neuroblastoma cell lines exhibit differing cell cycle and differentiation responses to ASCL1 overexpression. By comparing genome-wide ASCL1 chromatin binding, transcriptional changes, and protein-protein interactions, we found that ASCL1 binds more extensively to neuronal proteins in a cell line that is more susceptible to ASCL1-driven differentiation, but associates with cell cycle regulators in less responsive cells. We show that CDK2-Cyclin A2 bind ASCL1 in less responsive cells, with CDK-mediated phosphorylation of ASCL1 limiting the ability of ASCL1 to drive differentiation. Our study reveals that context-dependent interactions of ASCL1 with protein partners on the chromatin control its ability to re-engage a differentiation program in neuroblastoma.

## Introduction

Neuroblastoma is a cancer of the peripheral nervous system that accounts for approximately 15% of all paediatric cancer-related deaths^1^. It arises due to a differentiation block in the sympathoadrenal progenitor cells within the neural crest^2^. In normal development, sympathoadrenal precursor cells give rise to varied cell types, such as sympathetic neurons, Schwann cells and adrenal chromaffin cells^2^. Neuroblastoma can arise at different stages of sympathoadrenal development, resulting in a highly heterogeneous disease^3^.

The differentiation block and pro-proliferative state of neuroblastoma is thought to be epigenetically maintained; two oncogenic networks, or core regulatory circuits (CRC) of transcription factors that drive an adrenergic (ADRN) or a mesenchymal (MES) subset have been identified^4,5^. While ADRN cells are more committed to the adrenergic cell differentiation pathway and MES cells resemble a more undifferentiated phenotype^4,5^, heterogeneity is found even within each subtype.

This biological heterogeneity translates to a variety of clinical presentations, with a positive correlation between the degree of differentiation and long-term survival^6,7^. Notably, a subset of low-risk neuroblastoma regress spontaneously^8^, thought to result from re-engagement of a latent differentiation programme. In contrast, high-risk neuroblastoma, accounting for more than half of cases and frequently associated with *MYCN* amplification, is generally less differentiated and is associated with a worse prognosis and increased relapse rate^1,9,10^.

Given the close correlation between differentiation potential and favourable prognosis, differentiation therapies are a promising therapeutic approach in neuroblastoma^6,7,11,12^. Indeed, a differentiating agent, retinoic acid, is already employed in standard therapeutic regimes^13^. However, its efficacy is limited^14–16^; in order to develop more effective differentiation treatments, there is a vital need to understand the molecular mechanisms controlling the proliferation/differentiation balance in neuroblastoma.

Achaete-scute homolog 1 (ASCL1) is a basic helix-loop-helix (bHLH) transcription factor with a crucial role in balancing proliferation and differentiation in normal development, cancer and cell reprogramming. Acting as a pioneer transcription factor and a master regulator of neurogenesis^17,18^, ASCL1 directly binds closed chromatin, coordinating cell fate decisions^17–19^ and controlling the balance between proliferation and differentiation in varied contexts. For instance, ASCL1 loss- and gain-of-function studies show premature cell cycle withdrawal in neural progenitor cells and increased neuronal differentiation, respectively, in different *in vivo* and *in vitro* developmental models^18,20–22^. In the peripheral nervous system, ASCL1 is expressed transiently in the bridge population of the foetal adrenal medulla identified in a developmental trajectory, which connects Schwann cell precursors (SCPs) and progenitors that give rise to chromaffin cells or neuroblasts^23–25^. This suggests that ASCL1 may play key roles in sympathoadrenal cell fate decisions in normal development and tumorigenesis.

Consistent with its dual function, ASCL1 is reported to have context-dependent roles across different tumours, such as neuroblastoma^24,26^, small cell lung cancer^27,28^ and glioblastoma^29,30^, by promoting tumorigenesis or cell-cycle exit and differentiation. In neuroblastoma, ASCL1 is a component of the ADRN CRC that drives proliferation and promotes the expression of other CRC transcription factors^31,32^. In contrast, we have previously found that ASCL1 overexpression suppresses the proliferative programme and induces neuronal traits in the non-MYCN-amplified ALK-mutant neuroblastoma cell line SH-SY5Y^33,34^. Additionally, ASCL1 can reprogram non-neuronal cell types, such as fibroblasts, into neurons^35,36^. However, not all cells are responsive to ASCL1-induced reprogramming or differentiation. For example, keratinocytes fail to respond to ASCL1-induced reprogramming^36^. Similarly, MES-type neuroblastoma cells only partially respond to ASCL1 overexpression^37^. Despite its crucial role in balancing proliferation and differentiation, the regulation of ASCL1-induced differentiation in neuroblastoma is not fully understood, particularly in high-risk MYCN-amplified cases with poor prognoses.

Here, we explore the ability of ASCL1 to drive cell cycle exit and differentiation in MYCN-amplified neuroblastoma, the most common manifestation of high-risk disease. We find that two MYCN-amplified cell lines show different phenotypic responses to ASCL1 overexpression, reflected by differences in ASCL1’s genome-wide chromatin binding and transcriptional target activation. To explore the role of protein partners in this differential regulation we quantitatively compare the protein-protein interactors of ASCL1, finding that ASCL1 associates more with neuronal associated proteins in cells that respond to ASCL1 overexpression by undergoing differentiation. In contrast, ASCL1 associates more with positive cell-cycle regulators in cells that are less responsive to ASCL1-induced differentiation. Mutation of the Cy/RXL motif in ASCL1, responsible for association with cyclin proteins, enabled the less responsive cells to differentiate more effectively after ASCL1 over-expression. This study demonstrates that the context-dependent association of ASCL1 with factors including positive cell cycle regulators on chromatin plays an important role in controlling the proliferation/differentiation switch in neuroblastoma.

## Results

### ASCL1 overexpression alters neuronal differentiation and reduces proliferation in two MYCN-amplified neuroblastoma cell lines

We investigated the ability of increased ASCL1 levels to drive neuronal differentiation of the MYCN-amplified IMR-32 and SK-N-BE(2)C neuroblastoma cell lines, where endogenous ASCL1 expression is compatible with a proliferative progenitor identity^31^, by providing exogenous ASCL1. A doxycycline (dox)-inducible system was used for controlled overexpression of ASCL1, resulting in cell lines hereafter referred to as IMR^ASCL1^ and BE^ASCL1^ (Figure 1A). Western blotting confirmed elevated ASCL1 protein at similar levels in both cell lines upon dox treatment (Figures 1B and S1A), allowing direct comparison of the effects of overexpression.

**Figure 1.**
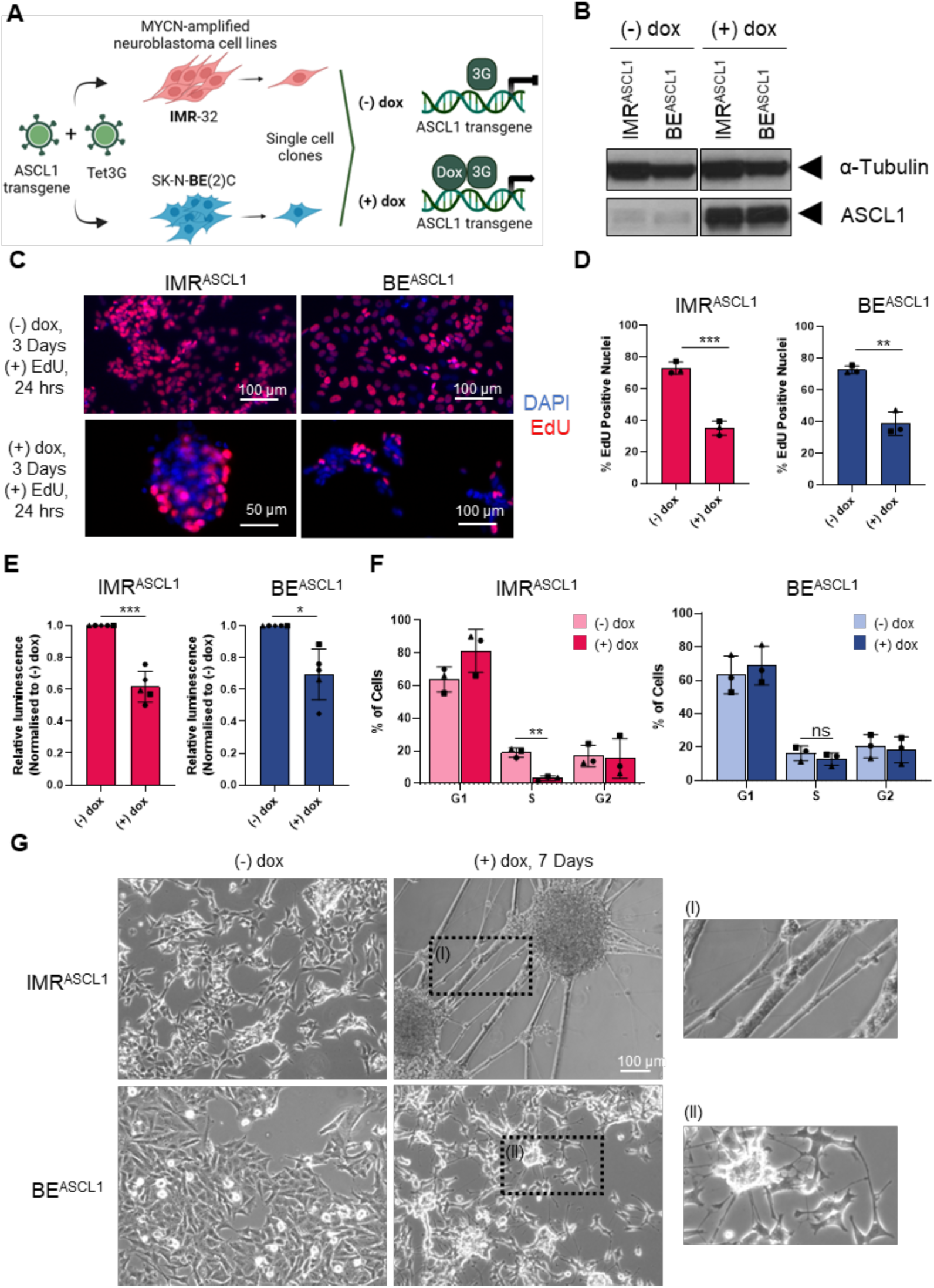
ASCL1 overexpression in MYCN-amplified neuroblastoma drives neuronal differentiation and reduced proliferation. **(A)** Schematic of the experimental system. Illustrates IMR-32 and SK-N-BE(2)C MYCN-amplified cell lines transduced with lentiviral vector encoding ASCL1 under the control of a Tet-On responsive element and a lentiviral vector encoding Tet3G. The two cell lines were single-cell sorted and expanded into clonal cell lines (IMR^ASCL1^ and BE^ASCL1^). Doxycycline (dox) treatment induces expression of ASCL1 transgene. **(B)** Representative western blot analysis of ASCL1 protein levels in IMR^ASCL1^ and BE^ASCL1^ cell lines treated with 1 µg/mL doxycycline (dox) for 24 hours. α-Tubulin is used as the loading control. n=3. **(C)** Representative images of EdU staining in IMR^ASCL1^ and BE^ASCL1^ cells after a 3 day 1 µg/mL dox treatment and an EdU pulse in the final 24 hours. DAPI in blue and EdU in red. Scale bars: 100 or 50 μm, as labelled. n=3. **(D)** Analysis of the percentage (%) of EdU-positive cells in (C). Mean ± SD, n = 3. Unpaired t test, ** p <= 0.01, *** p <= 0.001. **(E)** Cell number analysis using CellTiter-Glo assay 3 days after 1 µg/mL dox treatment of IMR^ASCL1^ and BE^ASCL1^ cells. For each cell line, luminescence values were normalised to untreated control (equals 1). Mean +/− SD, n = 5. One sample t test on dox-treated samples against the value 1 (normalised untreated control), *, p <= 0.05; ***, p <= 0.001. **(F)** Flow cytometry analysis of the percentage of cells in G1, S or G2 phases of the cell cycle, before and after ASCL1 overexpression for 24 hours with 1 µg/mL dox treatment, in IMR^ASCL1^ and BE^ASCL1^ cells. Mean +/− SD, n = 3. Unpaired t test, ns non-significant, ** p <= 0.01. **(G)** Representative phase-contrast images of IMR^ASCL1^ and BE^ASCL1^ cells 7 days after treatment with 1 µg/mL of dox. Treatment was not renewed for the duration of the 7 days. Scale bars: 100 μm. n=3. See also Figure S1.

First, we assessed the effect of ASCL1 overexpression on cell proliferation and morphology. IMR^ASCL1^ and BE^ASCL1^ showed a reduction in proliferation after 3 days of ASCL1 overexpression (Fig 1C-1D), as measured by EdU incorporation and CellTiter-Glo assays. Approximately 35.1% and 38.7% of cells were positive for EdU in dox-treated IMR^ASCL1^ and BE^ASCL1^, respectively, compared to 72.9% and 72.7% of untreated cells (Figure 1C-1D). The relative cell number was also significantly reduced in both cell lines upon ASCL1 overexpression (Fig 1E). Overexpression of ASCL1 significantly reduced the S phase population of IMR^ASCL1^ after 24 hours, while the cell cycle distribution of BE^ASCL1^ was unaltered (Figure 1F), indicating differences in the cell cycle response to ASCL1 overexpression.

After 7 days of dox treatment, IMR^ASCL1^ underwent striking morphological changes associated with neuronal differentiation (Figure 1G); cells formed large aggregates connected by long neurites, as has been previously observed to be associated with neuronal differentiation^33^. In contrast, BE^ASCL1^ showed more limited morphological changes, with modest cell clustering and shorter neurites (Figure 1G). Nevertheless, both cell lines upregulated βIII-tubulin, a marker of neural differentiation (Figure S1B). We conclude that, while ASCL1 can induce some morphological and cell cycle changes associated with neuronal differentiation in both MYCN-amplified neuroblastoma cell lines, the two cell lines respond differently to increased ASCL1 expression.

**Supplementary Figure 1 (Related to Figure 2):**
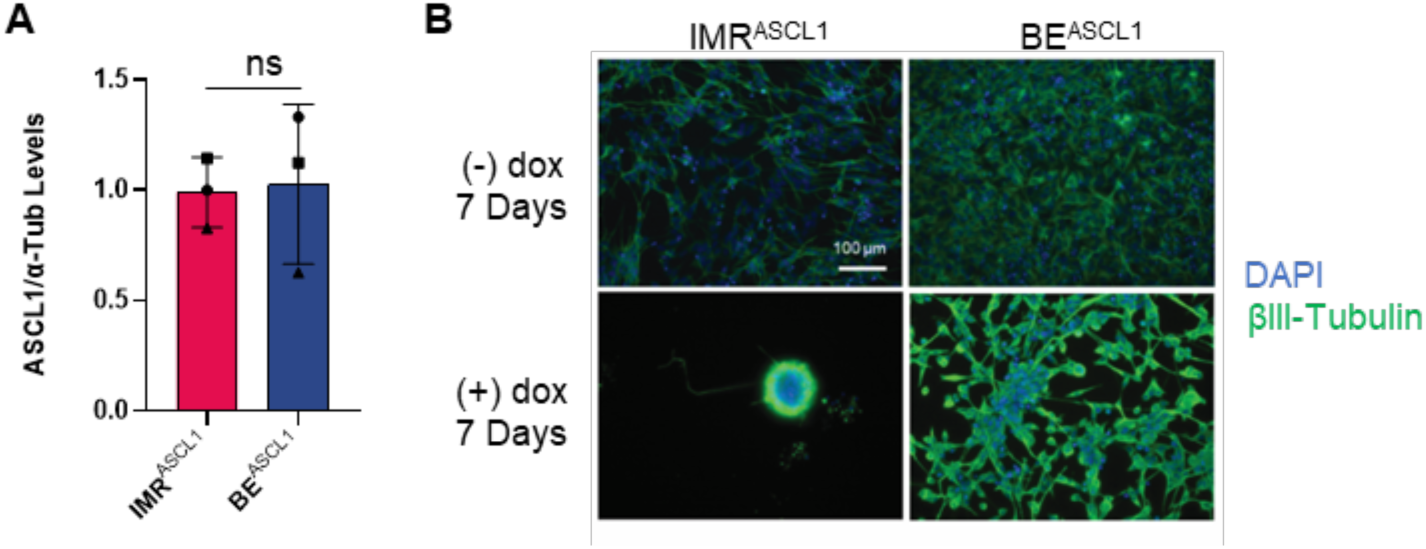
ASCL1 overexpression in MYCN-amplified neuroblastoma drives neuronal differentiation and reduced proliferation. **(A)** Quantification of ASCL1 levels from three biological repeats of western blots (Figure 1B). Plotted values represent the relative ASCL1 protein levels of IMR^ASCL1^ and BE^ASCL1^ cell lines treated with 1 µg/mL dox for 24 hours, normalised to α-Tubulin loading control. Mean ± SD, n = 3. Unpaired t test, ns non-significant. **(B)** Representative image of immunofluorescent staining using anti-βIII-tubulin antibody and DAPI, of IMR^ASCL1^ and BE^ASCL1^ cells treated with 1 µg/mL dox for 7 days. Scale bar: 100 μm. n=3.

### ASCL1 overexpression promotes greater transcriptional changes associated with increased neuronal differentiation and reduced proliferation in IMR-32 than in SK-N-BE(2)C cells

To characterise this differential response further, bulk RNA sequencing was performed on IMR^ASCL1^ and BE^ASCL1^ cells treated with dox for 24 hours. For each cell line, differential expression analysis was performed using DESeq2^38^, comparing dox-treated and untreated samples to ascertain the transcriptional changes induced by ASCL1 overexpression in the two contexts. In IMR^ASCL1^ cells, ASCL1 overexpression led to downregulation of 3,546 genes and upregulation of 4,169 genes (Figure S2, left-hand side), while in BE^ASCL1^ cells, there were a smaller number of transcriptional changes (1,730 genes downregulated and 1,967 genes upregulated, Figure S2, right-hand side).

To probe the function of the differentially regulated genes unique to each cell line and those common in both, we selected genes whose expression changed significantly in at least one cell line and clustered them into 8 groups, based on whether the genes were activated or repressed in the two lines. This revealed a complex pattern of differential gene expression in IMR^ASCL1^ and BE^ASCL1^ in response to ASCL1 (Figure 2A). Gene ontology analysis found that both cell lines upregulated known ASCL1 targets, such as the Notch signalling pathway and genes associated with positive regulation of neurogenesis and nervous system development (Figure 2B), consistent with the neuronal morphology induced by ASCL1. However, regulators of nervous system development were also downregulated in both cell lines, suggesting ASCL1 may exert complex control over different components of neuronal pathways in a context-dependent manner (Figure 2B). Furthermore, in line with the suppression of S phase only in IMR^ASCL1^, ASCL1 downregulated DNA replication genes specifically in IMR^ASCL1^ and not in BE^ASCL1^. In addition, upregulation of genes associated with synaptic processes found uniquely in IMR^ASCL1^ (Figure 2B) is consistent with their more mature neuronal phenotype compared to _BE_^ASCL1^.

**Figure 2.**
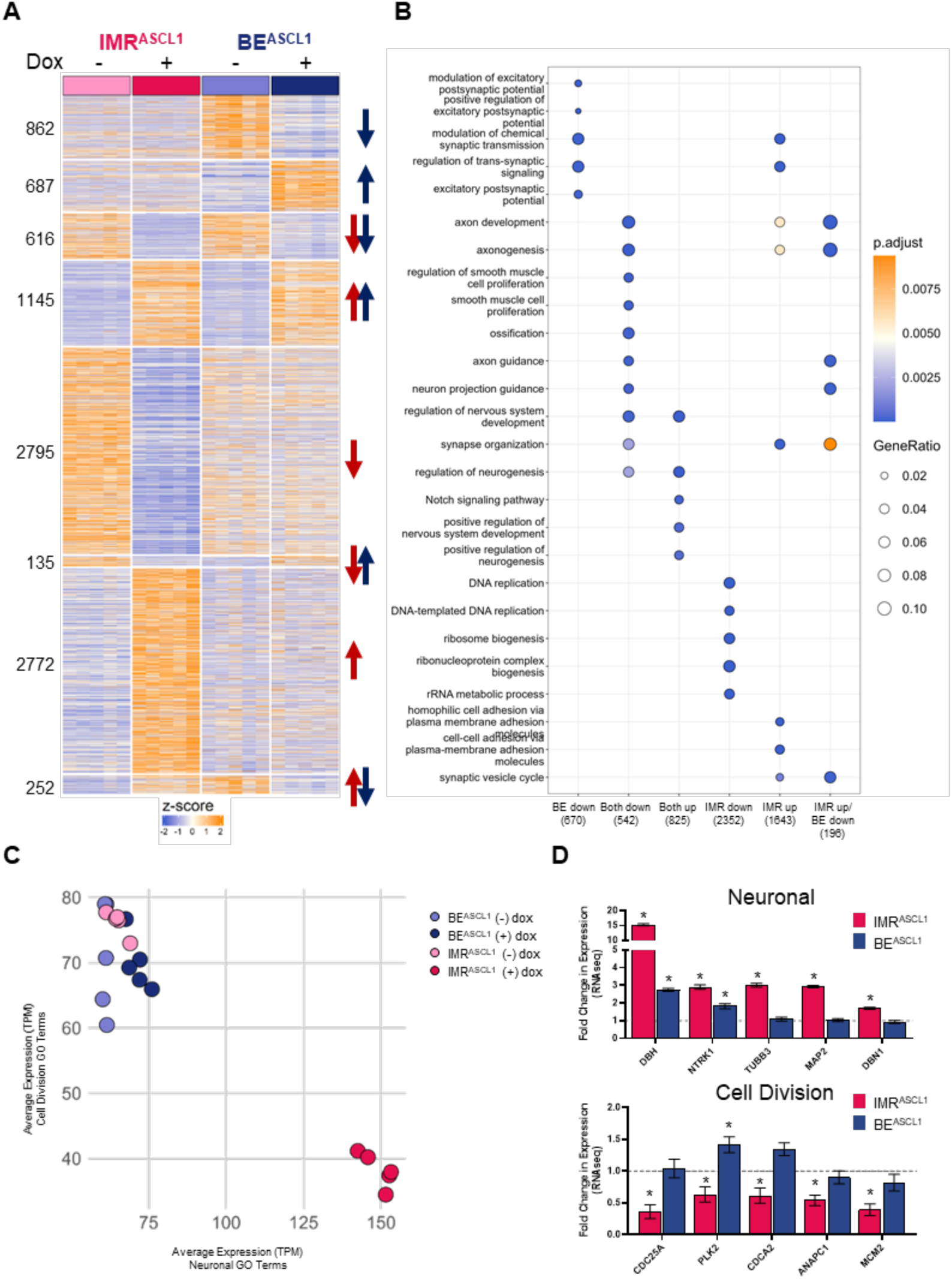
ASCL1 overexpression induces more transcriptional changes associated with neuronal differentiation in the IMR^ASCL1^ than BE^ASCL1^ cell line. **(A)** Heatmap of differentially expressed genes in either IMR^ASCL1^ or BE^ASCL1^ alone, or both cell lines upon upregulation of ASCL1 with 1 µg/mL doxycycline for 24 hours. A z-score has been applied to counts per million (CPM) normalised RNA sequencing counts across both cell lines. Differentially expressed genes were selected as follows: Downregulated, log2 fold change <-0.5 and adjusted p value < 0.05; Upregulated, log2 fold change >0.5 and adjusted p value < 0.05. Arrows indicate the direction of gene expression change for the relevant cell line. Blue arrows represent significant changes in BE^ASCL1^ and red arrows represent significant changes in IMR^ASCL1^. The number of genes in each group is noted to the left of the plot. **(B)** Gene ontology analysis of biological processes for the groups of genes represented in (A) using clusterProfiler. **(C)** Dot plot illustrating the average expression of neuronal (x axis) or cell division (y axis) gene signatures in IMR^ASCL1^ and BE^ASCL1^ cell lines untreated ((-) dox) or treated with 1 µg/mL dox ((+) dox) for 24 hours. Mean of Transcript counts per million (TPM) normalised RNA sequencing counts were used to plot average expression of neuronal or cell division gene signatures. Gene signatures were derived based on gene ontology groups associated with these processes enriched upon ASCL1 overexpression in either cell line. **(D)** Fold change in expression of dox-treated over untreated control samples for each cell line (IMR^ASCL1^ and BE^ASCL1^), for neuronal (top) or cell division (down) related genes. Error bars represent standard error values for log2 fold change from the DESeq2 analysis output. * represents significantly changing (log2 fold change > |0.5| and adjusted p value < 0.05) gene expression in dox-treated versus untreated, based on DESeq2 analysis. See also Figure S2.

Next, we specifically assessed expression of neuronal or cell division-related genes, by defining neuronal and cell division gene signatures. When comparing the average expression of each gene signature, we found that untreated IMR^ASCL1^ and BE^ASCL1^ samples clustered together (Figure 2C), indicating similar basal expression of both gene signatures in the absence of upregulated ASCL1. However, while dox-treated IMR^ASCL1^ demonstrated increased neuronal and reduced cell division gene signatures clustering away from the untreated samples, dox-treated BE^ASCL1^ remained clustered with their untreated counterparts (Figure 2C). Consistent with this, we saw greater ASCL1-mediated activation of differentiation targets such as *DBH*, *NTRK1*, *TUBB3*, *MAP2* and *DBN1* in IMR^ASCL1^ than BE^ASCL1^ (Figure 2D).

IMR^ASCL1^ also showed clear suppression of positive regulators of cell division such as *CDC25A, PLK2, CDCA2, ANAPC1* and *MCM2*, which was not observed in BE^ASCL1^ (Figure 2D). These data indicate that ASCL1 overexpression drives more pronounced transcriptional changes associated with neuronal differentiation and suppressed cell division in IMR-32 than SK-N-BE(2)C cells, reflecting the morphological and cell proliferation analyses described above (Figure 1).

**Supplementary Figure 2 (Related to Figure 2):**
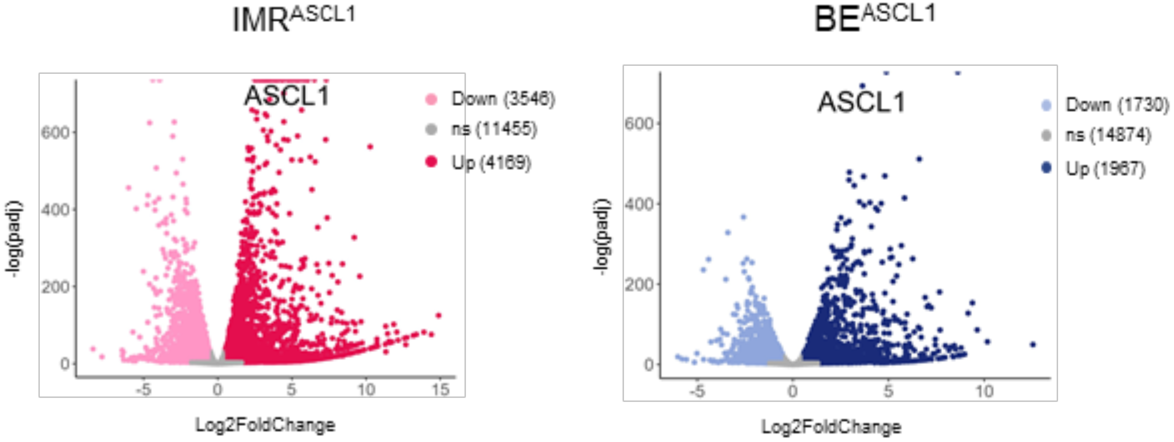
ASCL1 overexpression induces more transcriptional changes associated with neuronal differentiation in the IMR^ASCL1^ than BE^ASCL1^ cell line. Volcano plots illustrating the change in gene expression in dox-treated compared with untreated IMR^ASCL1^ (left) and BE^ASCL1^ (right) cell lines, based on DESeq2 analysis of RNA sequencing data. Significantly downregulated (down, log2 fold change < −0.5 and adjusted p value < 0.05), not significantly changing (ns, log2 fold change < |0.5| and/or adjusted p value > 0.05), and significantly upregulated (up, log2 fold change > 0.5 and adjusted p value < 0.05) genes are highlighted in pink, grey and red, respectively, for IMR^ASCL1^ cells, and in light blue, grey and dark blue, respectively, for BE^ASCL1^ cells. ASCL1 is labelled amongst the upregulated genes.

### ASCL1 binding and gene activation differs significantly between IMR^ASCL1^ and BE^ASCL1^

Next, we investigated why similar levels of ASCL1 drive more pronounced neuronal differentiation in IMR^ASCL1^ compared to BE^ASCL1^. Since target specificity of transcription factor binding can directly influence its function and, consequently, the transcriptional profile of the cells, we performed ASCL1 chromatin immunoprecipitation sequencing (ChIP-seq) on IMR^ASCL1^ and BE^ASCL1^ cells treated with doxycycline for 24 hours, using a well-validated anti-ASCL1 antibody^33^. This revealed that ASCL1 bound at more genomic sites in IMR^ASCL1^ than in BE^ASCL1^ (Figures 3A and S3A). Comparison of bound peaks between the two lines revealed 12,640 sites bound similarly by ASCL1 in both cell lines (Common peaks, Figure 3A). 13,458 sites showed increased ASCL1 binding in BE^ASCL1^ compared to IMR^ASCL1^ (BE enriched peaks, Figure 3A). However, the majority of the ASCL1-bound sites were significantly enriched in IMR^ASCL1^ versus BE^ASCL1^ (55,363 IMR enriched peaks, Figure 3A), indicating greater binding of ASCL1 to the chromatin in IMR^ASCL1^.

**Figure 3.**
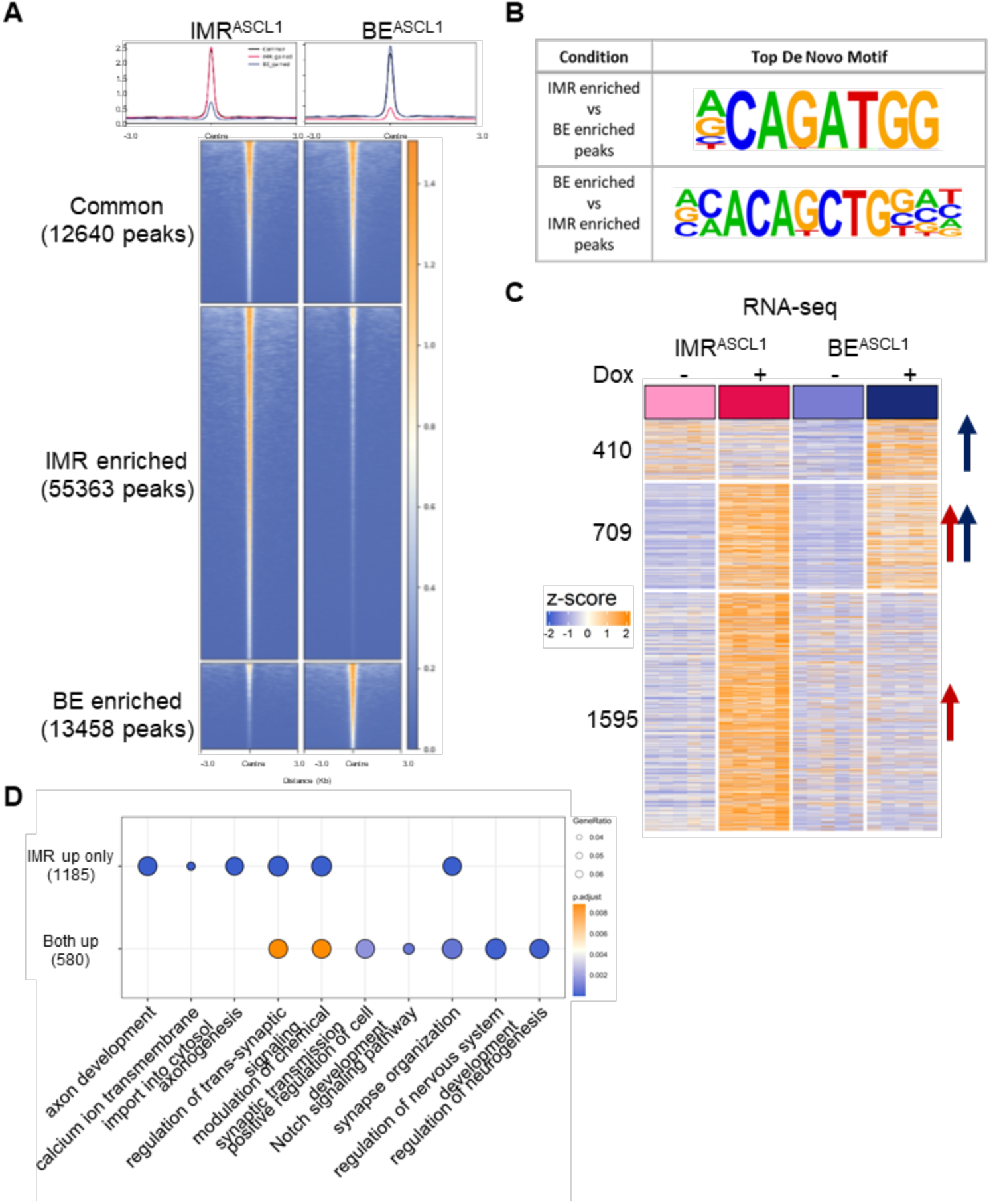
ASCL1 shows enhanced genome-wide chromatin binding in IMR^ASCL1^ versus BE^ASCL1^ cell line upon its overexpression. **(A)** Average profile (top) and aligned heatmap (bottom) illustration ASCL1 binding at Common (False discovery rate (FDR) > 0.1), IMR enriched (log2 fold change < −1 and FDR < 0.01), and BE enriched (log2 fold change >1 and FDR < 0.01) peaks. Data shown ± 3 kb from the Centre. Number of peaks in each category is shown. n = 4. **(B)** Top enriched *de novo* motif in IMR enriched (log2 fold change < −1 and FDR < 0.01) ASCL1 peaks compared to BE enriched (log2 fold change >1 and FDR < 0.01) ASCL1 peaks (top), and BE enriched ASCL1 peaks compared to IMR enriched ASCL1 peaks (bottom). Differential motif discovery analysis was performed using HOMER. **(C)** Heatmap showing relative gene expression of genes proximal (up to 50kb) to IMR enriched peaks in (A) and upregulated by ASCL1 overexpression, either in IMR^ASCL1^ or BE^ASCL1^ alone or both cell lines treated with dox for 24 hours. A z-score has been applied to counts per million (CPM) normalised RNA sequencing counts, across both cell lines. n=5. Arrows indicate the direction of gene expression change for the relevant cell line. Blue arrows represent significant changes in BE^ASCL1^ and red arrows represent significant changes in IMR^ASCL1^. The number of genes in each group is noted to the left of the plot. **(D)** Gene ontology analysis of biological processes comparing the 3 gene groups in (C). Gene ontology analysis performed using clusterProfiler. Genes upregulated in BE^ASCL1^ only did not show any enrichment. See also Figure S3.

Motif discovery analysis on ASCL1-bound sites found, as expected, an E-box DNA binding motif (5’-CANNTG-3’) to be the most significantly enriched motif in both cell lines^34^ (Figure S3B). Notably, however, a direct comparison of IMR enriched and BE enriched ASCL1 peaks revealed a bias towards a non-consensus E-box motif in IMR^ASCL1^ (5’-CAGATG-3’, Figure 3B). The overall enhanced chromatin binding and increased degeneracy of ASCL1 in IMR^ASCL1^ may reflect a more permissive cellular environment for ASCL1 binding that could contribute to enhanced cell cycle inhibition and differentiation compared to BE^ASCL1^.

To understand the functional relevance of the increased ASCL1 binding in IMR^ASCL1^ compared to BE^ASCL1^, we integrated ASCL1 ChIP and RNA sequencing data. We studied expression changes of genes proximal to IMR enriched ASCL1 peaks. As previous studies suggest that ASCL1 is a direct transcriptional activator but only an indirect transcriptional repressor^17,34^, we focused on bound and upregulated genes only, a set of 2,714 genes across the two cell lines. Analyses confirmed that the increased ASCL1 binding at these sites resulted in upregulation of a greater number of genes in IMR^ASCL1^ (2,304) compared to BE^ASCL1^ (1,119, Figure 3C). The group of genes uniquely upregulated in IMR^ASCL1^ resulting from enhanced ASCL1 binding, may reflect the greater phenotypic responsiveness of IMR^ASCL1^ cells to ASCL1-mediated neuronal differentiation, and this group is indeed enriched for genes with synaptic and axon development functions (Figure 3D). This highlights that the lower affinity binding sites enriched in IMR^ASCL1^ become functional and drive new gene signatures.

In conclusion, while the two MYCN-amplified neuroblastoma cell lines could be thought of as offering similar environments for activation of ASCL1 targets when it is expressed at similar levels, our results found many differences in ASCL1 binding and activity that correlate with the different phenotypic responses observed. This demonstrates the importance of context-dependent activity even for ASCL1, a potent pioneer factor^36,39^. We next wanted to understand why ASCL1 binding and transcriptional activity differed so significantly in these two MYCN-amplified neuroblastoma contexts.

**Supplementary Figure 3 (Related to Figure 3):**
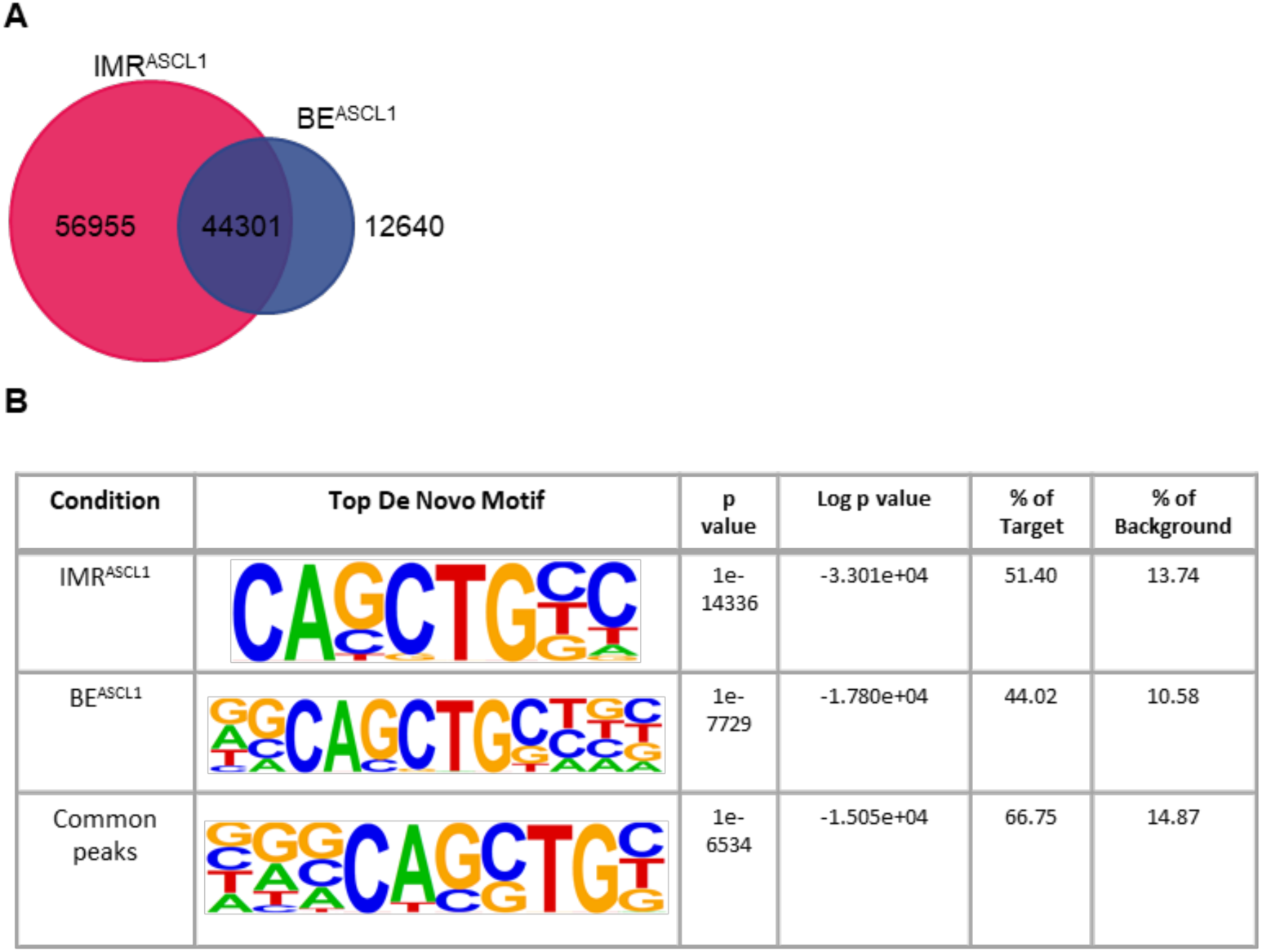
ASCL1 shows enhanced genome-wide chromatin binding in IMR^ASCL1^ versus BE^ASCL1^ cell line upon its overexpression. **(A)** Venn diagram comparing ASCL1 ChIP sequencing consensus peaks in IMR^ASCL1^ (left-hand side, red) and BE^ASCL1^ (right-hand side, blue) cell lines treated with 1 µg/mL dox for 24 hours. Numbers of unique and overlapping ASCL1 peaks are stated. n = 4. **(B)** Top enriched *de novo* motif in ASCL1 consensus peaks of IMR^ASCL1^ or BE^ASCL1^ dox-treated cell lines, as well as of sites Common to both cell lines according to DiffBind analysis (FDR>0.1). Differential motif discovery analysis was performed using HOMER, using a GC-matched background.

### The ASCL1 interactome is enriched for positive regulators of neuronal differentiation in IMR^ASCL1^ and cell cycle components in BE^ASCL1^

ASCL1 interactors play a crucial role in the function and regulation of ASCL1 during normal development and adult stem cell maintenance^40–42^, and potentially in balancing proliferation and differentiation in cancer. However, the identities and functions of ASCL1 interactors in neuroblastoma are largely unknown. To identify and quantitatively compare the chromatin-associated co-factors of ASCL1, Quantitative Multiplexed Rapid Immunoprecipitation Mass spectrometry of Endogenous proteins (qPLEX-RIME)^43,44^ was performed; this is a technique combining Chromatin Immunoprecipitation (ChIP), isobaric peptide labelling using Tandem Mass Tags, and peptide identification using MS3 mass spectrometry, allowing for quantitative comparison of chromatin-associated protein interactors of ASCL1 between the cell lines.

Prior to analysis, quality controls were performed. ASCL1 protein was successfully recovered with the 6 unique peptides identified covering approximately 18.64% of the ASCL1 sequence, mainly corresponding to the basic helix-loop-helix (bHLH) region of the protein (Figure S4A), ASCL1 immunoprecipitation was consistent across the cell lines (Figure S4B), and in each cell line, the enriched proteins in the ASCL1 immunoprecipitation compared to the IgG control included ASCL1 itself and the well-described ASCL1 binding partners TCF3, TCF4, TCF12 and HUWE1 (Figure S4C)

On investigating quantitative differences in the ASCL1 interactome, of the 912 ASCL1-associated proteins identified, 38 were significantly enriched in IMR^ASCL1^ and 56 significantly enriched in BE^ASCL1^ (Figure 4A). Gene ontology analysis was then conducted on the subsets of common interacting proteins, IMR^ASCL1^ enriched or BE^ASCL1^ enriched proteins. Protein partners found to bind ASCL1 similarly in both cell lines (818 proteins) were enriched for chromatin remodellers and mRNA processing proteins (Figure 4B-4C), likely reflecting ASCL1’s known function as a pioneer transcription factor. Amongst these were several SWI/SNF factors, previously shown to aid ASCL1-mediated neuronal differentiation in embryonic stem cells^39^, and histone modifiers such as P300, HDAC1, KDM1A and DNMT1 (Figure 4B-4C). Notably, ASCL1 bound more to proteins linked to regulation of neuron differentiation in IMR^ASCL1^ than BE^ASCL1^ (Figure 4B-4C). In contrast, in BE^ASCL1^, ASCL1 bound more to proteins associated with cyclin-dependent protein kinase (CDK) activity, such as the CDK2 and Cyclin A2 complex (Figure 4B-4C). In addition, we found other cell cycle associated proteins enriched in the BE^ASCL1^ ASCL1 interactome compared to that of IMR^ASCL1^, including SKP2 and PLK1 (Figure 4C).

**Figure 4.**
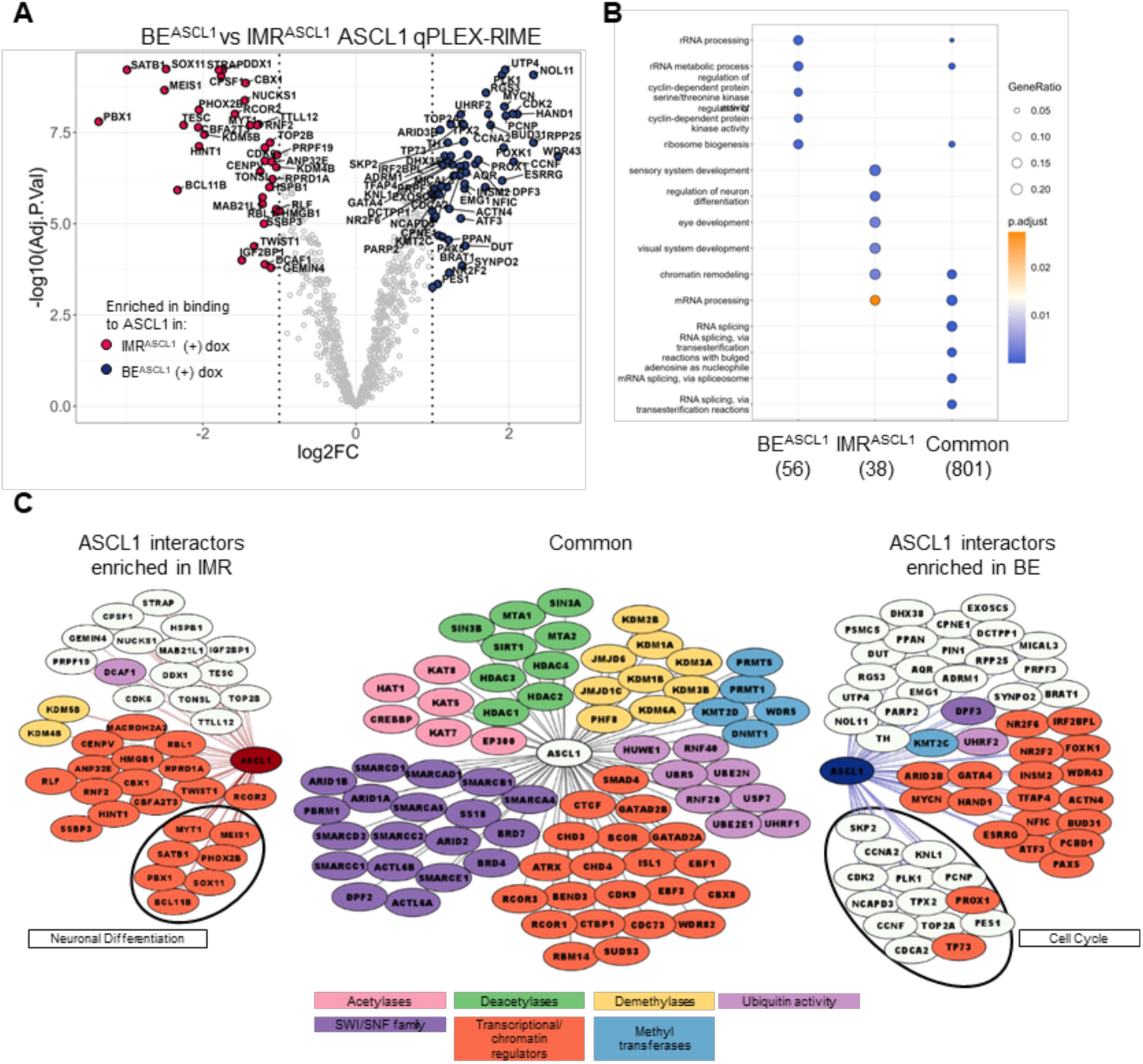
ASCL1 protein-protein interactome is enriched for regulators of neuronal differentiation in the IMR^ASCL1^ cell line and cell cycle components in the BE^ASCL1^ cell line. **(A)** Volcano plot of the quantitative analysis, by qPLEX-RIME, of proteins associated to ASCL1 in BE^ASCL1^ versus IMR^ASCL1^ cell lines, treated with 1 µg/mL doxycycline for 24 hours. Proteins enriched in the ASCL1 interactome in the IMR^ASCL1^ cell line are highlighted in red (log2 fold change < −1 and adjusted p value < 0.05) and those enriched in the ASCL1 interactome in the BE^ASCL1^ cell line are highlighted in blue (log2 fold change > 1 and adjusted p value < 0.05). n=5. **(B)** Ontology analysis of ASCL1 interactors enriched in the IMR^ASCL1^, neither, or the BE^ASCL1^ cell line from (A). **(C)** Cytoscape diagram of ASCL1 interactors. On the left-hand side are illustrated ASCL1 interactors enriched in the IMR^ASCL1^ cell line. In the centre are illustrated shortlisted ASCL1 interactors not enriched in either cell line. On the right-hand side ASCL1 interactors enriched in the BE^ASCL1^ cell line are shown. Known functions of these proteins are colour coded or highlighted. See also Figure S4.

Further, key proteins in neuroblastoma pathogenesis such as N-MYC, HAND1, GATA4, ARID3B and TFAP4 associated with ASCL1 more in BE^ASCL1^ compared to IMR^ASCL1^. N-MYC and HAND1 contribute to neuroblastoma pathogenesis as part of the ADRN CRC which maintains cells in a proliferative state^5^, while ARID3B and TFAP4 work cooperatively with N-MYC to drive an aggressive phenotype in neuroblastoma^45,46^. In contrast, a number of proteins important during normal neuronal development such as BCL11B^47^, PBX1^48^ and SATB1^49,50^ were associated with ASCL1 more in IMR^ASCL1^ (Figures 4A-4C). PBX1 was previously shown to be important in retinoic acid-induced differentiation of neuroblastoma cells as well as acting as a favourable prognostic biomarker^51^.

The enriched neuronal differentiation regulators in IMR^ASCL1^, and positive cell cycle regulators in BE^ASCL1^ indicate that proteins associating with ASCL1 may influence its ability to engage a more pro-differentiation or pro-proliferative transcriptional programme, depending on the cellular context.

**Supplementary Figure 4 (Related to Figure 4):**
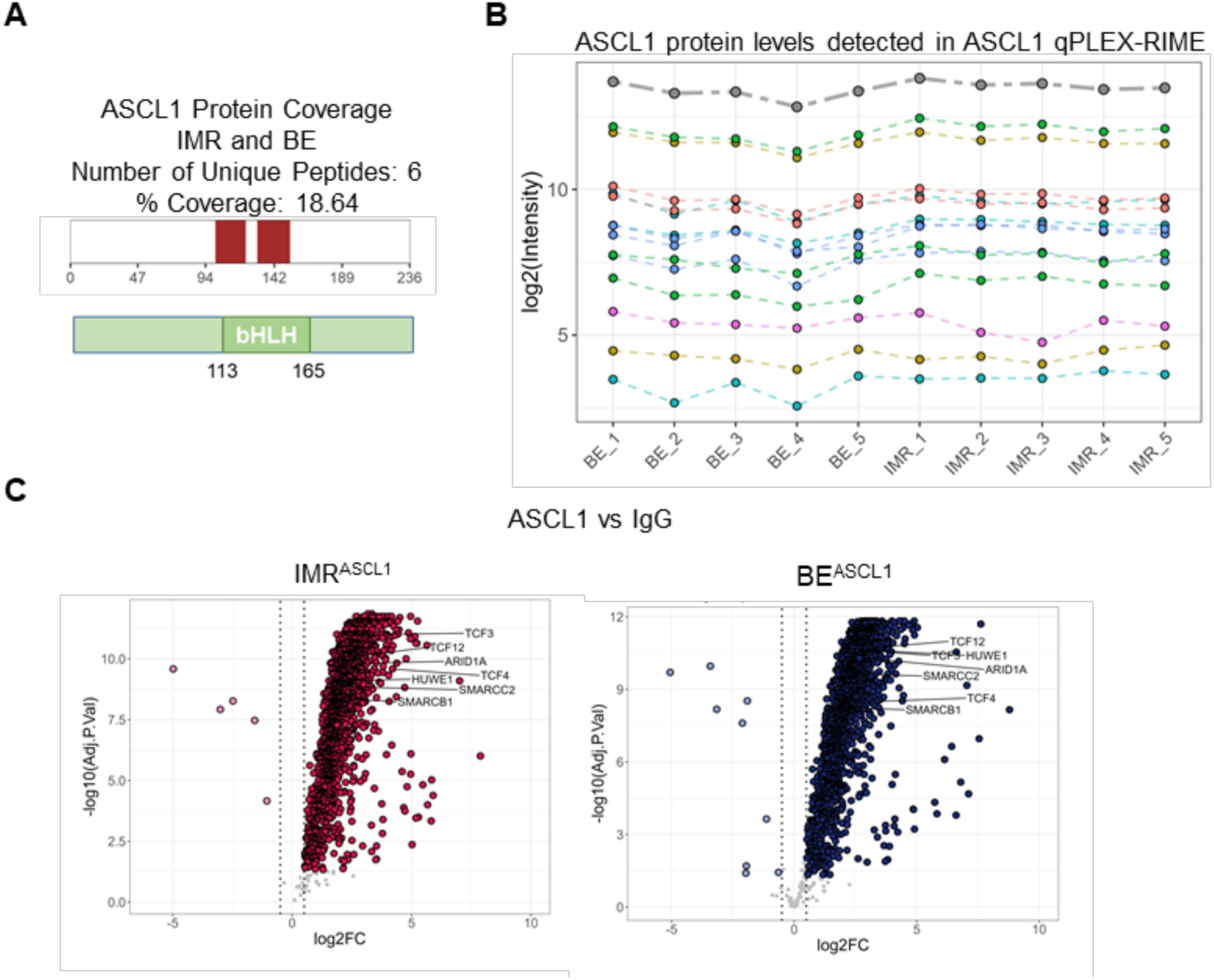
ASCL1 protein-protein interactome is enriched for regulators of neuronal differentiation in the IMR^ASCL1^ cell line and cell cycle components in the BE^ASCL1^ cell line. **(A)** Sequence coverage of the ASCL1 protein in the ASCL1 qPLEX-RIME experiment in IMR^ASCL1^ and BE^ASCL1^ cell lines overexpressing ASCL1, and schematic of its basic helix-loop-helix domain. n=5. **(B)** Log2 Intensity of ASCL1 peptides, in colour, and their sum, in grey, across 5 biological replicates of ASCL1 qPLEX-RIME in IMR^ASCL1^ and BE^ASCL1^ cell lines treated with 1 µg/mL doxycycline for 24 hours. **(C)** Volcano plots of ASCL1 vs IgG pull-downs in the qPLEX-RIME experiment in IMR^ASCL1^ (left-hand side) and BE^ASCL1^ (right-hand side) cell lines. Several known ASCL1 interactors from literature are labelled. Proteins were considered enriched in the ASCL1 pull-down if adjusted p value < 0.05 and log_2_ fold change > 0.5.

### CDK1 and CDK2 phosphorylate ASCL1 affecting its chromatin binding

Given that the cell cycle machinery has been reported to affect the function of bHLH master regulators during normal development^22,52–54^, we further interrogated the interaction of ASCL1 with Cyclins and CDKs detected in the ASCL1 qPLEX-RIME (Figure 4). In both cell lines Cyclins and CDKs were found to associate with ASCL1; However, CDK2, Cyclin A2 and Cyclin F were significantly enriched in the ASCL1 interactome in BE^ASCL1^ compared to IMR^ASCL1^ (Figure 5A). Although below the fold change threshold for significance, other CDKs, such as CDK1 and CDK9, also showed a tendency to bind to ASCL1 more in BE^ASCL1^ than IMR^ASCL1^ (Figure 5A). Quantitative measurement of the total proteome of IMR^ASCL1^ and BE^ASCL1^ demonstrated that total protein levels of CDKs and Cyclins enriched in in the BE^ASCL1^ ASCL1 interactome were very similar in the two cell lines (Figure 5B). We therefore conclude the differences in CDK/Cyclin association with ASCL1 are due to enhanced binding in BE^ASCL1^ rather than differences in overall CDK/Cyclin protein availability.

**Figure 5.**
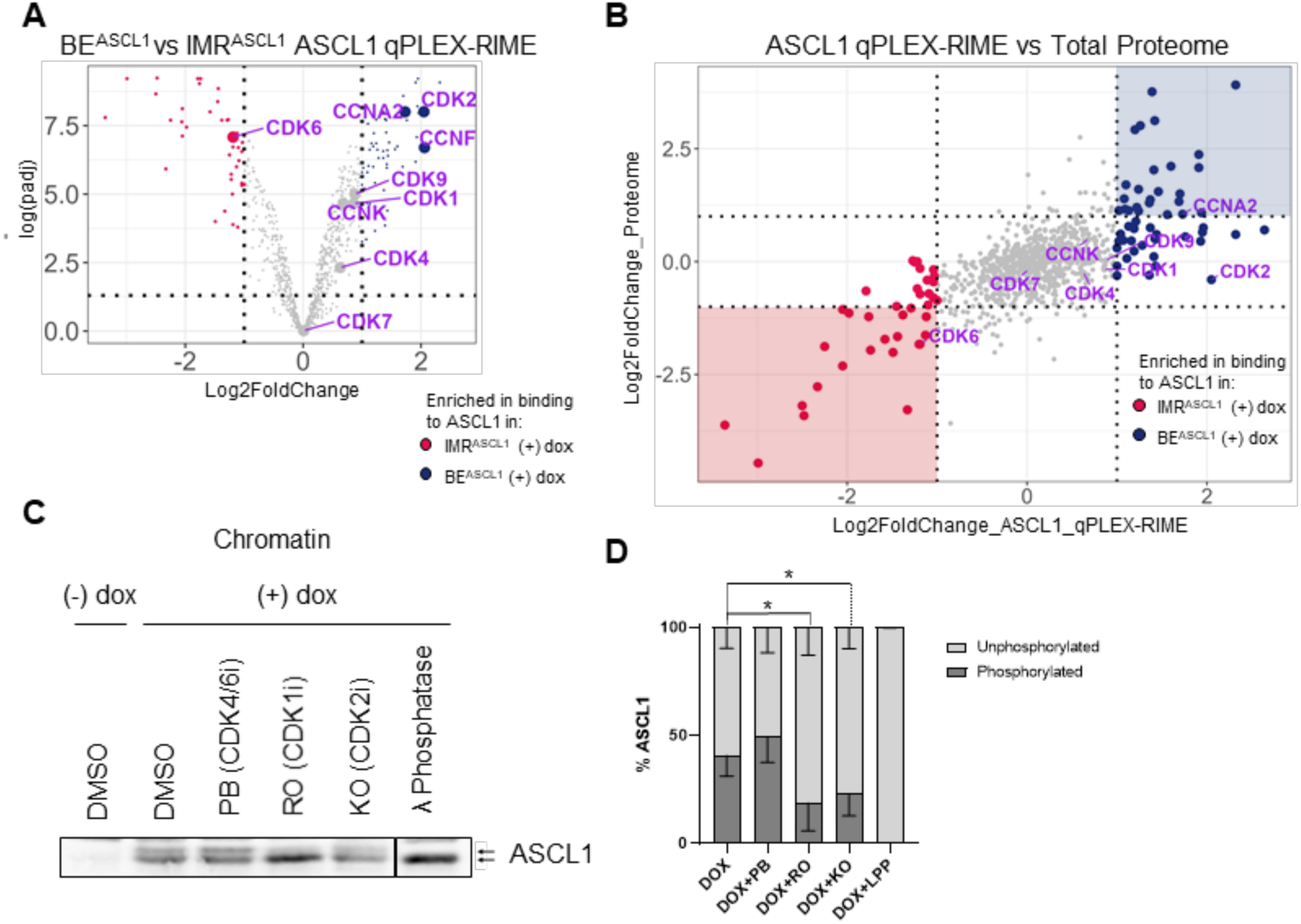
CDK1 and CDK2 associate with and phosphorylate ASCL1, affecting its binding to the chromatin in BE^ASCL1^ cells. **(A)** Volcano plot of quantitative ASCL1 interactome analysis (qPLEX-RIME) between IMR^ASCL1^ and BE^ASCL1^ cell lines treated with 1 µg/mL dox for 24 hours, highlighting CDK and Cyclin proteins detected. In blue, are highlighted proteins associating with ASCL1 more in BE^ASCL1^ cells compared to IMR^ASCL1^, and in red, are highlighted proteins associating with ASCL1 more in the IMR^ASCL1^ cells compared to BE^ASCL1^. Proteins are considered enriched when |log2 fold change| >1 and adjusted p value < 0.05. **(B)** Dot plot comparing Total Proteome and ASCL1 qPLEX-RIME in BE^ASCL1^ and IMR^ASCL1^ cell lines treated with 1 µg/mL dox for 24 hours. In red are highlighted proteins enriched in binding to ASCL1 in IMR^ASCL1^ cells, and in blue the proteins enriched in binding to ASCL1 in BE^ASCL1^ cells (qPLEX-RIME). Labelled are all CDKs and Cyclins detected in the ASCL1 qPLEX-RIME. **(C)** Representative western blot analysis of ASCL1 protein in the chromatin fraction of BE^ASCL1^ cells. The cells were left untreated or treated with dox for 24 hours, in combination with small molecule CDK inhibitors of CDK4/6 (5 µM of Palbociclib, PB), CDK1 (16 µM of RO3360, RO) or CDK2 (8 µM of KO, KO). Λ Protein Phosphatase treatment was used as control for un-phosphorylated ASCL1. Two western blot lanes, between KO and λ Protein Phosphatase, were cut out the blot in PowerPoint. n=4. **(D)** Quantification of western blot analysis in (C), shown as percentage of phosphorylated and un-phosphorylated ASCL1. Mean ± SD, n=4. Unpaired t test, * p <= 0.05. See also Figure S5.

CDK-mediated phosphorylation of ASCL1 has been shown to affect its activity in developing frog embryos and mammalian cells^22,24,55^. Consequently, we further explored the possible significance of differential regulation of ASCL1 by the cell cycle machinery in BE^ASCL1^ and IMR^ASCL1^. *In vitro* kinase assays show that CDK1/Cyclin B, CDK2/Cyclin A and CDK2/Cyclin E complexes can phosphorylate human ASCL1 (Figure S5). To investigate how this phosphorylation could affect ASCL1 chromatin binding in neuroblastoma cells, we inhibited CDK4/6, CDK1 or CDK2 activity in BE^ASCL1^ with small molecule CDK inhibitors Palbociclib (PB)^56,57^, RO-3306 (RO)^58^, or KO3861 (KO)^59^, respectively, concomitant with ASCL1 induction for 24 hours. Assessment of the phospho-status of ASCL1 bound to chromatin showed that CDK1 and CDK2 inhibition increased the ratio of unphosphorylated to phosphorylated chromatin-bound ASCL1 (Figure 5C-5D), while CDK4/6 inhibition had no effect.

**Supplementary Figure 5 (Related to Figure 5):**
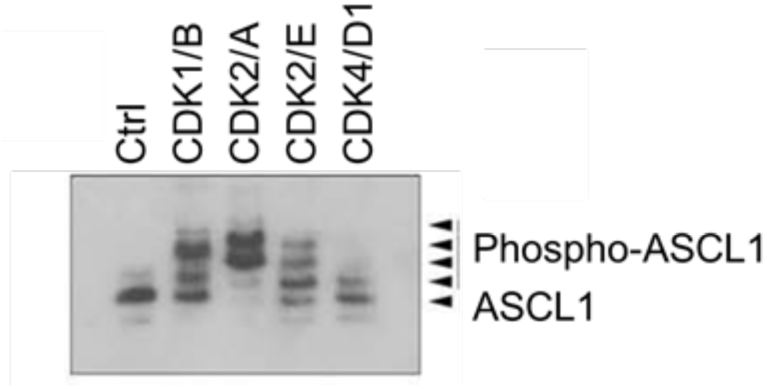
CDK1 and CDK2 associate with and phosphorylate ASCL1, affecting its binding to the chromatin in BE^ASCL1^ cells. Phos-tag western blot analysis of *in vitro* kinase assay with translated HA-tagged human wild-type ASCL1 and human recombinant Cyclin-CDK complexes, such as CyclinB-CDK1, CyclinA-CDK2, CyclinE-CDK2 or CDK4-CyclinD1. n=3.

### Disruption of CDK/Cyclin-mediated regulation of ASCL1 improves ASCL1-driven neuronal differentiation

Finally, we wanted to assess the functional role of the identified ASCL1-CDK/Cyclin interaction in regulating ASCL1-induced differentiation, hypothesising that ASCL1 association with and phosphorylation by CDK/Cyclin complexes on chromatin directly suppresses its ability to drive differentiation in BE cells. For this purpose, we first mutated a conserved Cy/RXL motif (KRRL(100-103)AAAA, Figure 6A) in ASCL1, identified in a diverse range of proteins including CDK inhibitors to mediate interactions with Cyclins^60^. The KRRL motif we identified and the surrounding sequence in ASCL1 was a close match to part of the P21 CDK inhibitor protein (Figure S6A), shown to facilitate interactions and inhibition of CDK/Cyclin complexes^61^. Secondly, we mutated all potential serine-proline CDK phosphorylation sites in ASCL1 to generate a phospho-mutant (S-A) ASCL1, shown to drive differentiation and maturation of neurons in embryonic development, in reprogramming and in non-MYCN-amplified ALK-mutant neuroblastoma cells^22,24,33,34^.

**Figure 6.**
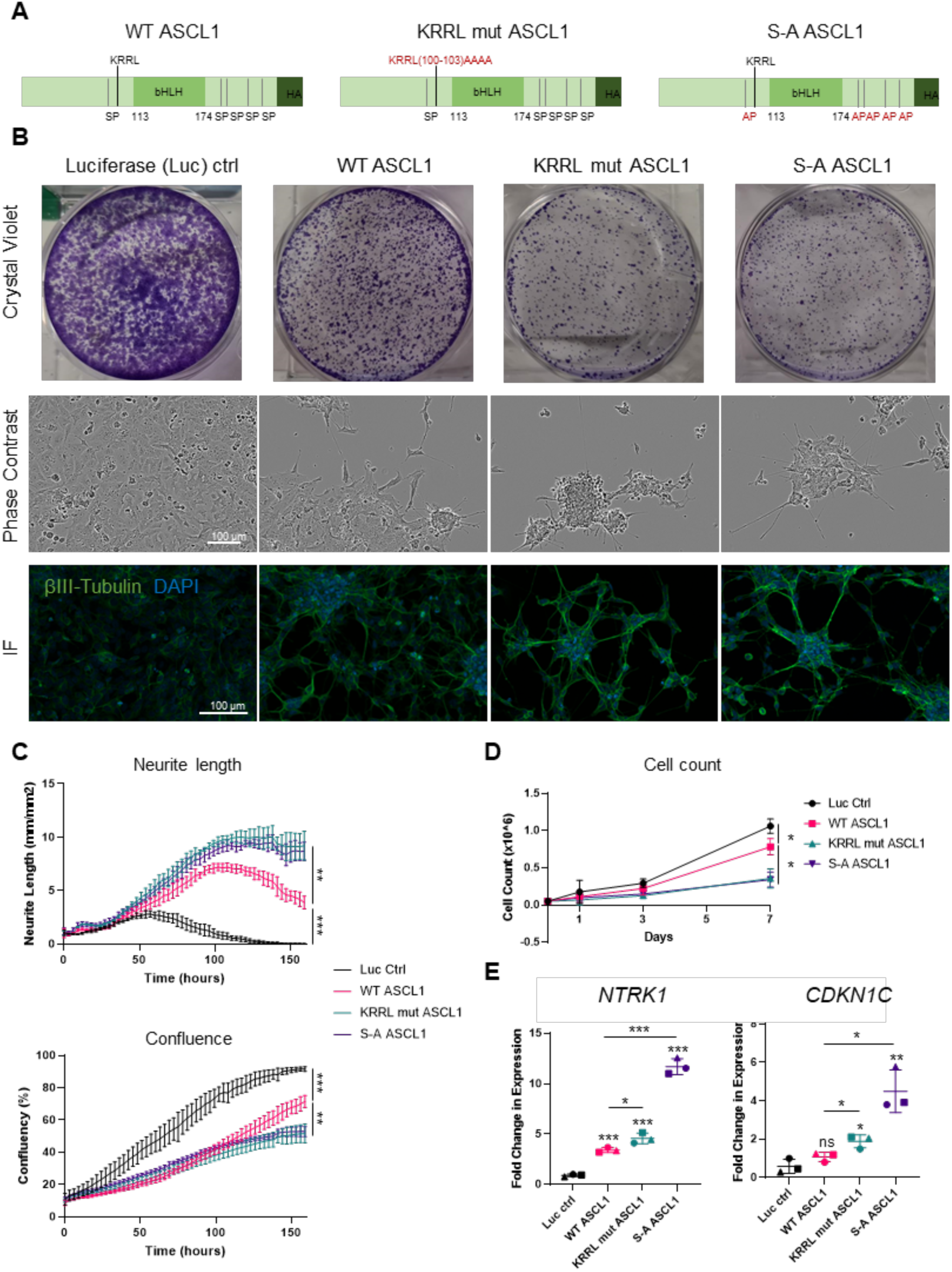
Overexpression of Cy/RXL mutant or phospho-mutant ASCL1 improves neuronal differentiation of BE cells over wild-type ASCL1. **(A)** Schematic of the wild-type (WT), KRRL mutant (mut) and phospho-mutant (S-A) ASCL1 proteins that were overexpressed in the BE cell line using a lentiviral and TET-On inducible expression system. The figure illustrates the KRRL and SP sites mutated in red, and the presence of the C-terminus HA-tag. **(B)** Representative crystal violet staining, phase contrast and immunofluorescent (IF) staining of βIII-tubulin images of BE cells transduced with a luciferase control, wild-type ASCL1, KRRL mutant ASCL1 or S-A ASCL1. Cells were treated with 1 µg/mL doxycycline for 7 days. n=3. **(C)** Quantification of average Neurite Length (top) and Confluence (bottom) of BE cells overexpressing a Luciferase control (Luc ctrl), wild-type (WT) ASCL1, KRRL mutant (mut) ASCL1 or phospho-mutant (S-A) ASCL1 over the course of 7 days post 1 µg/mL doxycycline treatment. Imagining and quantification was performed on an IncuCyte®. Mean ± SD, n=3. Unpaired t test, ** p <= 0.01, *** p <= 0.001. **(D)** Cell count of BE cells overexpressing a Luciferase control (Luc ctrl), wild-type (WT) ASCL1, KRRL mutant (mut) ASCL1 or phospho-mutant (S-A) ASCL1 on days 1, 3 and 7 post 1 µg/mL doxycycline treatment. Mean ± SD, n=3. Unpaired t test on day 7 samples, * p <= 0.05. **(E)** Quantitative PCR analysis of the relative expression of *NTRK1* or *CDKN1C* in BE cells overexpressing a Luciferase control (Luc ctrl), wild-type (WT) ASCL1, KRRL mutant (mut) ASCL1 or phospho-mutant (S-A) ASCL1. Cells were treated with 1 µg/mL doxycycline for 7 days. Mean ± SD, n=3. Unpaired t test, ns non-significant, * p <= 0.05, ** p <= 0.01, *** p <= 0.001. * directly above data points represent comparisons to the Luciferase control sample. See also Figure S6.

Using the same lentiviral and dox-inducible expression system used to overexpress wild-type ASCL1 (Figures 1-5), wild-type (WT), KRRL(100-103)AAAA mutant (KRRL mut), phospho-mutant (S-A) ASCL1, or a luciferase control (Luc ctrl) were overexpressed in BE cells. ASCL1 protein levels were similar in WT and KRRL mutant overexpressing BE cells after 24 hours, with higher S-A ASCL1 protein being detected (Figure S6B), possibly related to somewhat enhanced stability of phospho-mutant ASCL1^22,55^. Co-immunoprecipitation of HA-tagged ASCL1 confirmed disruption of the interaction between ASCL1 and Cyclin A2 in BE cells overexpressing KRRL mutated ASCL1 (Figure S6C).

Strikingly, overexpression of KRRL mutant or S-A ASCL1 further reduced proliferation, and improved the differentiated morphology of BE cells compared to WT ASCL1 (Figure 6B), as seen in whole-well crystal violet staining, phase-contrast imaging and immunofluorescent staining of neuronal marker βIII-Tubulin (Figure 6B). Increased neurite length and reduced cell confluence of BE cells overexpressing the KRRL mutant or S-A ASCL1 compared to WT ASCL1 was observed over 7 days after ASCL1 overexpression (Figure 6C). Cell counting was also performed, with BE cells overexpressing KRRL mutant or S-A ASCL1 showing significantly fewer cells compared to WT ASCL1 at day 7 (Figure 6D). Furthermore, compared to the Luciferase control cells, expression of *NTRK1,* a neuronal marker and marker of favourable prognosis in neuroblastoma^62^, was somewhat elevated by WT ASCL1 overexpression, but was greater in KRRL mutant ASCL1-overexpressing cells and greater still after S-A ASCL1 overexpression (Figure 6E). The CDK inhibitor *CDKN1C* (P57), a member of the Cip/Kip family of CDK inhibitors that has the ability to inhibit the activity of all cell cycle regulating CDK/Cyclin complexes^63,64^, was also upregulated by KRRL mutant and S-A ASCL1 but not by WT ASCL1 in BE cells. These data support an enhanced ability of KRRL mutant or S-A ASCL1 to drive neuronal differentiation and inhibit proliferation in BE cells compared to WT ASCL1, and further demonstrates the role of CDK-dependent regulation of ASCL1 activity by direct association of Cyclin/CDK complexes to ASCL1 on chromatin.

**Supplementary Figure 6 (Related to Figure 6):**
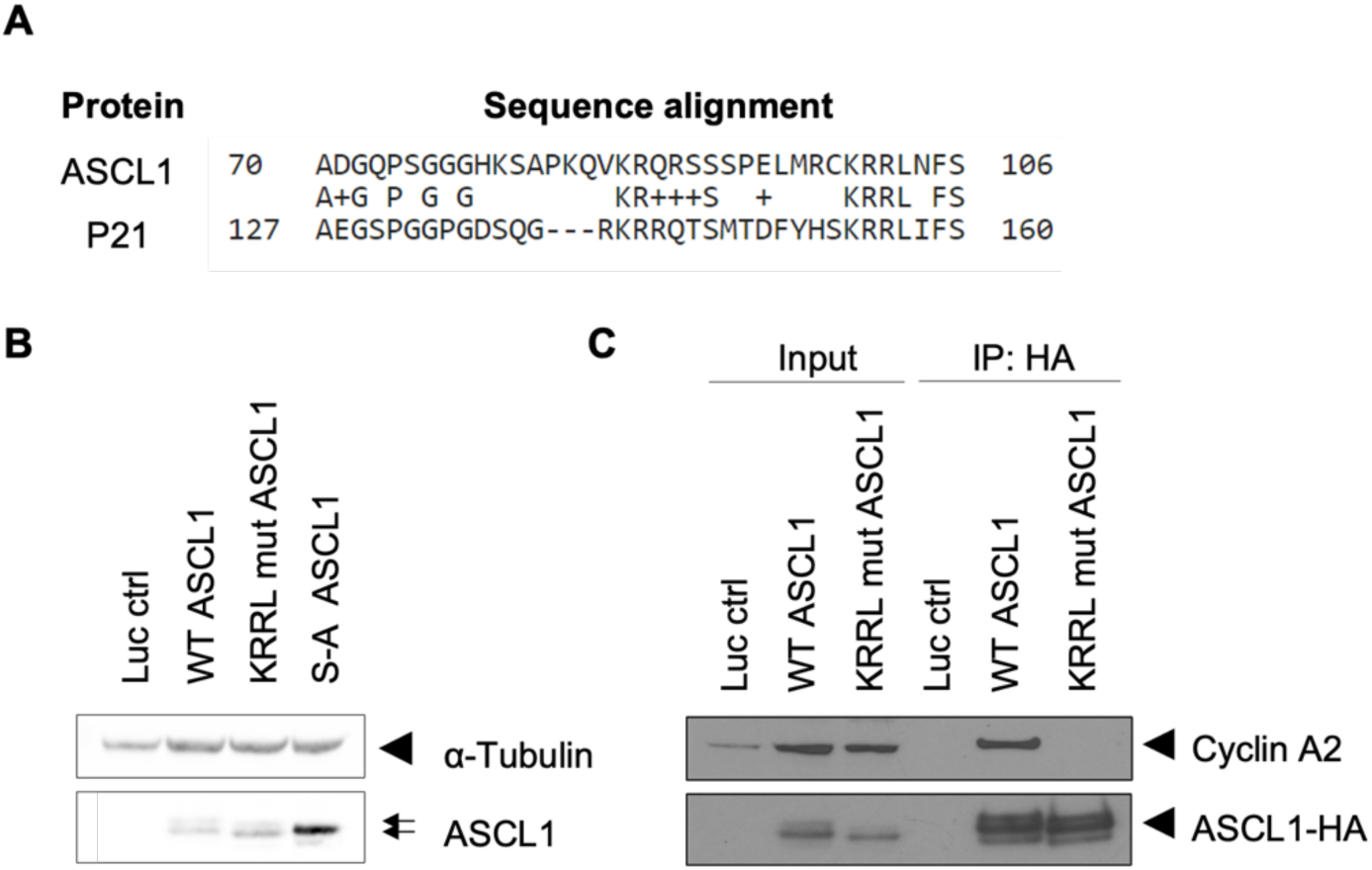
Overexpression of Cy/RXL mutant or phospho-mutant ASCL1 improves neuronal differentiation of BE cells over wild-type ASCL1. **(A)** Protein BLAST of ASCL1 and CDK inhibitor P21, illustrating the alignment between the two protein sequences, including the Cy/KRRL motif. **(B)** Western blot analysis of ASCL1 protein levels in BE cells overexpressing a Luciferase control (Luc ctrl), wild-type (WT) ASCL1, KRRL mutant (mut) ASCL1, or phospho-mutant (S-A) ASCL1. Cells were treated with 1 µg/mL doxycycline for 24 hours. α-Tubulin was used as a loading control. n=3. **(C)** Western blot analysis of co-immunoprecipitation (co-IP) of HA tagged ASCL1 and Cyclin A2 in BE cells overexpressing Luc ctrl, WT ASCL1, or KRRL mutant ASCL1. Cells were treated with 1 µg/mL doxycycline for 24 hours prior to HA co-IP. Western blot was probed for Cyclin A2 and HA (ASCL1). n=2.

## Discussion

Differentiation therapies are a promising therapeutic approach for the childhood cancer neuroblastoma^57^, but a lack of understanding of the switch from proliferation to differentiation in neuroblastoma hinders progress. A potent proneural factor, ASCL1, has been harnessed in the field of neuronal reprogramming, with different cell types showing varied susceptibility to ASCL1-induced neuronal differentiation^33–37,65,66^. What mediates this varied responsiveness to ASCL1 is still not fully understood, and understanding the mechanisms underlying the differentiation potential of neuroblastoma cells is crucial to the development of novel therapeutics.

In this study, we investigated the ability of ASCL1 to drive neuronal differentiation of adrenergic MYCN-amplified neuroblastoma cells, associated with high-risk disease. We found that, while ASCL1 overexpression led to enhanced neuronal characteristics in both MYCN-amplified IMR-32 and SK-N-BE(2)C cell lines, IMR^ASCL1^ showed a more robust neuronal differentiation morphology and transcriptional profile (Figures 1-3). This differential response highlights the heterogeneity in neuroblastoma and the importance of the cellular context for the responsiveness to differentiating cues.

RNA-seq and ChIP-seq analyses revealed increased ASCL1 binding and transcriptional activity at specific neuronal differentiation-associated sites in IMR-32 compared to SK-N-BE(2)C (Figure 3). This could arise due to differences in the epigenetic landscape of the two cell lines; for example, differing epigenetic marks could favour or limit ASCL1’s ability to associate to the chromatin in distinct sites. The E-box motif degeneracy we observed in ASCL1 binding sites in IMR^ASCL1^ compared to BE^ASCL1^ (Figure 3 and 3S) further supports a more permissive environment for ASCL1 to access and associate with lower affinity sites in the former. Indeed, ChIP-seq data from neuronal precursor cells and mouse embryonic fibroblasts overexpressing ASCL1 showed increased ASCL1 binding degeneracy in the neuronal context, indicating that more degenerate motifs are associated with a more permissive cellular environment for ASCL1 activity^34^. Therefore, the generation of *de novo* binding sites, particularly in genes associated with a neuronal phenotype, may be required for ASCL1 to exert its neurogenic role. Wapinski *et al.*^36^ suggested that ASCL1 does not reprogram keratinocytes due to lack of crucial histone marks (a trivalent epigenetic chromatin signature of H3K4me1, H3K27ac, and H3K9me3) at differentiation genes required for optimal ASCL1 binding and transcriptional activity, reinforcing the potential role of the epigenetic landscape in determining responsiveness.

To understand why ASCL1 binding and transcriptional activity differed so significantly in these superficially similar cellular contexts, we analysed co-factor availability, known to influence ASCL1 function in development^39,67–69^. qPLEX-RIME identified interactions of ASCL1 with chromatin remodellers in both cell lines, an interactome including proteins with acetylase, deacetylase, methylase and ubiquitin activity, many of which are previously undescribed ASCL1 interactors (Figure 4). ASCL1’s ability to induce chromatin remodelling at certain sites during neuronal differentiation of human stem cells is dependent on co-binding to DNA with SWI/SNF chromatin remodelling proteins^39^. We found SMARCB1 and other SWI/SNF proteins associated with ASCL1 in both IMR^ASCl1^ and BE^ASCL1^ cells (Figure 4), which may be important for the upregulation of common gene targets such as regulators of neurogenesis and Notch signalling genes.

Analysis of differences in the ASCL1 interactome between cell types (Figure 4) suggest ASCL1 may work in conjunction with other neuronal regulators to switch IMR^ASCL1^ cells from a proliferative to a differentiated phenotype, whereas in BE^ASCL1^ cells, lower association with pro-differentiation protein partners may limit its ability to drive neuronal maturation. Instead, proteins associated with ASCL1 in BE^ASCL1^ cells, namely positive cell cycle regulators including Cyclins and Cyclin-dependent kinases (CDKs), may actively promote a pro-proliferative and less differentiated phenotype. Indeed, the cell cycle and cell cycle machinery have been heavily implicated in regulation of differentiation^54,70^.

Post-translational modifications, particularly phosphorylation, often play crucial regulatory roles in control of transcription factor activity^22,24^. We have previously shown that ASCL1 is phosphorylated by CDKs in a Xenopus frog model of developmental neurogenesis, hindering its ability to drive neuronal differentiation^24,33^. Here, we find that CDK1 and CDK2 phosphorylate ASCL1 in neuroblastoma cells (Figures 5 and S5) and are associated with it on chromatin (Figures 4 and 5). Moreover, inhibition of CDK1 and CDK2 activity, but not CDK4/6, led to increased levels of unphosphorylated ASCL1 on chromatin in BE^ASCL1^ (Figure 5). These data are consistent with previous findings that support increased DNA binding and transcriptional activity of a phospho-mutant (S-A) ASCL1, with mutated serine-proline phosphorylation sites that prevents phosphorylation by CDKs, compared to wild-type (WT), in developmental and reprogramming models^22,24,33,34^. We also find overexpression of S-A ASCL1 in SK-N-BE(2)C cells resulted in reduced proliferation and increased morphological and marker-associated neuronal differentiation compared to WT ASCL1 (Figure 6), demonstrating that CDK-dependent phosphorylation normally limits its ability to engage a differentiation programme in these high risk MYCN-amplified tumour cells. Confirming a regulatory role for the direct association of CDK/Cyclin complexes with ASCL1 on chromatin, overexpression of a KRRL mutant form of ASCL1 that shows disrupted binding to Cyclin A2 (Figure S6) also improved ASCL1-induced neuronal differentiation of BE^ASCL1^ (Figure 6).

These data indicate that CDK1 and CDK2 bound to ASCL1 in BE^ASCL1^ may act directly via ASCL1 protein phosphorylation on chromatin to inhibit its ability to drive neuronal differentiation. The increased association of ASCL1 with CDK/Cyclin complexes in BE^ASCL1^ may also impact the phosphorylation of other proteins on chromatin where, in particular, CDK1 has been shown to phosphorylate many of its substrates, altering the epigenetic landscape of the cells^71^. Therefore, increased recruitment of CDKs to the chromatin by ASCL1 at critical targets could also indirectly alter the cells’ responsiveness to ASCL1-induced differentiation by affecting ASCL1 interactors and the wider chromatin landscape.

Our findings advance understanding of the control of neuronal differentiation and potential roadblocks in neuroblastoma, and in particular how the heterogeneity of neuroblastoma tumours may influence their responsiveness to potential differentiation therapies^57^. Understanding the roles of ASCL1 interactors in controlling activation of its diverse targets could help explain whether this key transcriptional regulator can support cell proliferation or promote differentiation of neuroblastoma in a context-dependent manner, allowing modulation of ASCL1-mediated differentiation to improve therapeutic options for this devastating disease.

## Materials and Methods

### Lead contact

Further information and requests for resources and reagents should be directed to and will be fulfilled by the Lead Contact, Professor Anna Philpott.

### Materials Availability

All custom plasmids and cell lines generated from this study will be available upon request.

### Data and code availability

Sequencing data has been deposited in the Gene Expression Omnibus data repository (GEO). RNAseq −/+ doxycycline of IMR^ASCL1^ and BE^ASCL1^ cell lines is included in the Series GSE213430. ASCL1 ChIP-seq + doxycycline of IMR^ASCL1^ and BE^ASCL1^ cell lines is included in the Series GSE213247. The mass spectrometry proteomics data have been deposited to the ProteomeXchange Consortium via the PRIDE^72^ partner repository with the dataset identifier PXD056448. Custom analysis code required to re-analyse the data in this study is deposited on Zenodo (doi: 10.5281/zenodo.13835001). Other original data is available upon request.

### Cell culture

IMR-32, SK-N-BE(2)C parental or engineered cell lines were cultured in DMEM-F12 media with L-glutamine (Gibco), further supplemented with 10% Foetal Bovine Serum (Sigma). Cell lines were verified by submitting genomic DNA for short tandem repeat sequencing and compared with data from the Cellosaurus database. All cell lines were confirmed to be Mycoplasma negative and were tested at a minimum of every 3 months. Cells were maintained at 37 °C and 5% CO_2_. To passage, seed or collect cells trypsin was used.

### Cell line treatments

#### Doxycycline Induction

Expression of ASCL1 in doxycycline-inducible cell lines was induced with 1 µg/mL.

#### Cyclin-Dependent Kinase (CDK) Inhibitor Treatment

Cells were treated with CDK4/6 inhibitor Palbociclib (PB, Selleckchem) at 5 µM, CDK1 inhibitor RO3306 (RO, Selleckchem) at 16 µM and CDK2 inhibitor KO3861 (KO, Selleckchem) at 8 µM for 24 hours in combination with 1 µg/mL doxycycline.

### Generation of Lentivirally Transduced Cell Lines

#### Construct preparation

A 3rd Generation Lenti-X vector (pLVX-TREG, Takara) was used as a backbone to clone untagged wild-type (WT) ASCL1 (previously described in ^33^), C-terminus HA-tagged WT ASCL1, C-terminus HA-tagged KRRL mutant ASCL1 or C-terminus HA-tagged S-A mutant ASCL1, using In-Fusion Cloning (Takara). A control plasmid expressing Luciferase (pLVX-TRE3G-Luc Control Vector) was purchased from Takara.

To generate C-terminus HA-tagged KRRL mutant ASCL1, In-Fusion Cloning mutagenesis was performed on WT ASCL1 in a pCS2+ vector. The primers 5’-GCGCAGCCGCGGCCAACTTCAGCGGCTTTGGC-3’ and 5’-TGGCCGCGGCTGCGCAGCGCATCAGTTCGGG-3’, along with PrimeStar GXL DNA polymerase (Takara) were used to mutate the 5’-AAACGCCGGCTC-3’ region of WT ASCL1 to 5’-GCAGCCGCGGCC-3’, creating a KRRL(100-103)AAAA mutation. A HA tag (YPYDVPDYA) was cloned on the C-terminus of ASCL1, using the same method.

C-terminus HA-tagged S-A mutant ASCL1 was generated by site directed mutagenesis using the QuikChange II XL Site-Directed Mutagenesis Kit (Agilent Technologies), as described in^33^.

#### Lentiviral preparation and transduction

Viral particles carrying a pLVX-TREG vector with the gene of interest or pLVX-CMV-Tet3G, encoding a Tet-On transactivator, were produced by transfecting the HEK 293T cell line with the ProFection Mammalian Tranfection System (Promega) together with a viral packaging mix containing PMD2G, PMLg, REV/PRSV, in a ratio of 6:3:4:2, respectively. HEK 293T cells were left to produce viral particles for ∼ 30 hours. Subsequently, cell media was collected and filtered to remove cell debris. The virus was concentrated overnight using Lenti-X concentrator (Takara) and collected by centrifugation at 4000 xg for 45 minutes at 4°C. Quantification was performed with the Lenti-XTM qRT-PCR Titration Kit (Takara).

IMR^ASCL1^ and BE^ASCL1^ cells were generated from IMR-32 and SK-N-BE(2)C cells, respectively, as previously described^33^. SK-N-BE(2)C cells expressing inducible WT ASCL1-HA, KRRL mutant ASCL1-HA, S-A ASCL1-HA or Luciferase control, were generated in a similar way but with a few differences. SK-N-BE(2)C cells were simultaneously transduced with viral particles carrying pLVX-CMV-Tet3G and viral particles carrying the gene of interest in pLVX-TREG, at a multiplicity of infection (MOI) of 10 each. The media containing viral particles was replaced with normal media after 24 hours. Following a further 24 hours, the cells expressing the vector were selected with 700 μg/mL G418 for 5 days, followed by 48 hour recovery and a second selection with 2 μg/mL Puromycin for 8 days. SK-N-BE(2)C WT ASCL1-HA, KRRL mutant ASCL1-HA, S-A ASCL1-HA and Luciferase control were used as pools.

### Flow Cytometry

#### Single-cell Fluorescence-activated Cell Sorting (FACS)

To isolate single cells into 96-well plates for clonal cell line generation, cells were dissociated, washed, resuspended in 1 mL of PBS and passed through a cell strainer. As a live/dead stain 5 µL of DR (ThermoFisher) was added to the cells. FACS performed on Aria-Fusion cell sorter with FACSDiva Software by the NIHR Cambridge BRC Cell Phenotyping Hub.

#### Cell-cycle Analysis using Flow Cytometry

Cells were seeded on a 10-cm dish and left untreated or treated with doxycycline for 24 hours. All incubations were performed at room temperature and on a gentle rotor. For cell collection and washes throughout the protocol, centrifugation was performed at 500 xg. Prior to cell fixation with 4% paraformaldehyde (PFA) for 10 minutes, cells were dissociated, collected and washed 3 times in 0.5% BSA in PBS. After fixation, cells were incubated with Hoersch (1:1000) in 0.5% BSA in PBS for 30 minutes. Another 3 gentle washes in PBS (with 0.5% BSA) were performed, cells resuspended in 1mL FACS buffer (5mM EDTA, 25mM HEPES buffer, 0.5% BSA in PBS, pH7) and passed through a cell strainer. Flow cytometry analysis was performed on BD LSRFortessa with FACSDiva software. FloJo was used for data analysis.

### Proliferation Analysis

#### EdU Staining

Cells were left untreated or induced to express ASCL1 with dox. On day 2, media with or without dox was renewed and supplemented with 10 µM EdU. Cells were incubated with EdU for 24 hours at normal culturing conditions and then fixed with 4% PFA for 10 minutes at room temperature. To stain for EdU, Click-iT EdU Alexa Fluor 647 assay kit (Thermo Fisher Scientific) was used according to manufacturer’s instructions. Nuclei were stained with DAPI. Imaging of stained cells was performed on Leica DMI6000 microscope using z-stacks. Maximum intensity z-projection was generated with FIJI software. Per biological condition, at least 100 nuclei were imaged and the percentage of EdU positive cells quantified using the CellProfiler software.

#### CellTiter-Glo® Luminescent Cell Viability Assay

For relative cell number quantification, CellTiter-Glo assay (Promega) was used as per manufacturer’s instructions. The CellTiter-Glo reagent was reconstituted and added to each 96-well plate at 100 µL/well, followed by vigorous shaking of the plate for 2 minutes. Lysed cells were incubated at room temperature without shaking for an additional 10 minutes, before quantifying luminescence on Spectra Max M5^e^ with softmax pro v.5.2 software. As a control for background signal, CellTitre-Glo reagent was added to wells containing only media.

#### Cell Counting

Cell counting was performed on Countess™ II FL Automated Cell Counter (Thermo Fisher Scientific), by diluting an aliquot of cells 1:1 in trypan blue stain. At least two technical repeats were averaged per cell count.

#### Cell confluence and neurite analysis using IncuCyte

Cells were plated on a 6 well-plate and live-cell imaged for 7 days post dox treatment on an IncuCyte® (Sartorius). Incucyte® Classic Confluence Analysis was used to study cell confluency over time, and Incucyte® Neurotrack Analysis Software Module was used to track neurite length.

#### Imaging

Cell imaging was carried out using Olympus IX51 microscope with a Leica DFC310 FX camera for phase-contrast images. Fluorescent microscopy was performed on Leica DMI6000 microscope. Imagine analysis performed on FIJI and CellProfiler.

#### ImmunoFluorescence Staining

Cells were left untreated or treated with dox for 7 days. After completion of treatment, cells were fixed with 4% PFA for 10 minutes, followed by 3 gentle washes with PBS. All incubations were performed at room temperature. Permeabilisation was achieved with an incubation with 0.2% Triton X-100 in PBS for 10 minutes. Blocking was performed in 3% BSA in PBS for 1 hour, followed by incubation with primary antibody against βIII-tubulin (1:2000) in antibody diluent (2% BSA, 0.2% Triton X-100 in PBS) for 1 hour. Cells were washes 3x in PBS and incubated with secondary antibody at 1:1000 in antibody diluent for 1 hour. After the incubation, another 3 washes with PBS were completed. Nuclei were stained with DAPI (1:5000) for 20 minutes. Cells were imaged on Leica DMI6000 microscope.

#### Crystal Violet Staining

Cells were seeded and treated on 6-well plates. At day 7 post dox treatment cells were fixed in 4% PFA for 10 minutes at room temperature. The cells were washed in PBS and stained with 0.5% aqueous crystal violet solution for 30 minutes at room temperature. Crystal violet was washed off gently with deionised water. Plates were allowed to dry before taking pictures.

#### Quantitative Total Proteome

For quantitative whole proteome analysis, IMR^ASCL1^ and BE^ASCL1^ cells were treated with 1 µg/mL doxycycline for 24 hours on 15-cm dishes. Cells were collected in PBS containing protease and phosphatase inhibitors (Roche), followed by centrifugation at 300 xg for 5 min. Pellets were washed 3 times in supplemented PBS. Similar sized pellets of at least 6 million cells were snap frozen on dry ice and submitted to the Proteomics facility at the Cancer Research UK Institute for Tandem Mass Tag (TMT) labelling and mass spectrometry analysis. Equal amounts of protein were used per sample after protein concentration was quantified using Bradford assay (BIO-RAD-Quick start). The subsequent sample processing and mass spectrometry analysis was performed as described in ^44^.

### Rapid Immunoprecipitation Mass-Spectrometry of Endogenous Proteins (RIME) and quantitative multiplexed RIME (qPLEX-RIME)

Cell lines were grown in 15-cm dishes in DMEM-F12 media supplemented with 10% TET-free FBS. Two 15-cm dishes of 80% confluent cells were cross-linked and collected for analysis per sample. An additional 15-cm dish was collected without cross-linking to quantify ASCL1 levels for normalisation. ASCL1 levels were assessed from the non-cross-linked samples via western blot, with ImageJ used for quantification. These quantified ASCL1 levels were then used to normalize the corresponding cross-linked samples before proceeding with immunoprecipitation.

Double cross-linking was performed using 2 mM DSG in PBS for 20 min, followed by 1% formaldehyde (FA) for 10 min. The reaction was quenched with 0.1 M glycine. For nuclear extraction, pelleted cells were resuspended in LB1 buffer and mixed by rotation for 10 minutes at 4 °C. Samples were subject to centrifugation (2,000 xg), and the resulting pellets were resuspended in LB2 buffer and rotated for 5 minutes at 4 °C. Another centrifugation step was performed, and pellets were resuspended in LB3 buffer. Chromatin was fragmented by sonication in a BioRuptor Plus sonication device (Diagenode) to produce DNA fragments of 100–1,000 base pairs, which were verified by gel electrophoresis on a 2% agarose gel.

Meanwhile, per sample, 50 µL of beads were conjugated to 5 µg of antibody overnight at 4°C. The bead-bound antibody was then added to the normalized and fragmented chromatin samples and rotated overnight at 4 °C. The following day, the beads were washed 10 times with 1 mL RIPA buffer and twice with 500 μL 100 mM AMBIC, on ice. AMBIC was prepared fresh each time. The tryptic digestion and mass spectrometry analysis were performed by the Proteomics facility at the Cancer Research UK Institute, as described in ^44^. Cytoscape was used to illustrate the interactors.

### Bioinformatics Analysis of Quantitative Total Proteome and ASCL1 qPLEX-RIME

The upstream analysis of collision induced dissociation (CID) tandem mass spectra were conducted by the Cancer Research UK Cambridge Institute Proteomic Facility as described in ^44^. The subsequent downstream analysis was performed using the qPLEXanalyzer^44,73^ Bioconductor package. The analysis was performed on unique peptides identified with high confidence (peptide FDR <1%).

For both Total Proteome and qPLEX-RIME, peptide intensities were normalised using group median scaling within each cell line. Control IgG samples were normalised separately, in the qPLEX-RIME. Proteins with adjusted p value < 0.01 and log_2_FC > 1 or log_2_FC < −1 were considered significantly different in protein levels (total proteome) between BE^ASCL1^ and IMR^ASCL1^ cell lines. Proteins with adjusted p value < 0.05 and log_2_FC > 1 or log_2_FC < −1 were considered significantly enriched in binding to ASCL1 (qPLEX-RIME) in BE^ASCL1^ or IMR^ASCL1^, respectively.

For the qPLEX-RIME experiments, missing values in IgG samples were replaced with the lowest value in the data, in this case 0.1. This ensured statistical analysis could be performed on peptides with low detection in unspecific IgG control. To filter out the non-specific interactors, proteins enriched in ASCL1 versus IgG pull-down samples were selected based on adjusted p value < 0.05 and log_2_FC > 0.5. After selecting the ‘specific’ interactors, the analysis was re-started with the refined list of proteins as stated above. The data were further filtered using the UniProt and CRAPome databases. Proteins known to localize to the nucleus, or those with no known localization according to UniProt, were retained. Conversely, proteins identified in over 50% of affinity purification experiments on human samples, as listed in the CRAPome database, were excluded from the analysis. The CRAPome database compiles proteins commonly detected in negative controls across various affinity purification mass spectrometry experiments.

### *In vitro* kinase assay

*In vitro* translated HA-tagged wild-type ASCL1 (SP6 TNT Quick Couple transcription kit, Promega) and 100nM of human recombinant Cyclin-CDK complexes, such as CyclinB-CDK1, CyclinA-CDK2, CyclinE-CDK2 or CDK4-CyclinD1, were incubated in the presence of ATP in NEB buffer (NEB) for 1 hour at 30°C. Resulting samples were analysed on a phos-tag western blot and probed using an anti-HA antibody.

### Quantitative RT-PCR (qRT-PCR)

Cells were lysed with RLT buffer and RNA extracted using the RNeasy Mini kit (Qiagen). RNA concentration was determined using a NanoDrop™ Spectrophotometer. Reverse transcription was performed using the QuantiTect Reverse Transcriptase Kit (Qiagen). Quantitative RT-PCR (qRT-PCR) was performed and on an Applied Biosystems StepOne™ Real-Time PCR system, using SYBR™ Green Master Mix (Thermo Scientific) and oligonucleotide primers (Sigma) specific for each gene of interest. The data were analysed using the ddCt method and are presented as fold changes in relation to one replicate in the control sample (Luciferase overexpressing BE cells), where this value equals one. Data presented as Mean +/− SD.

### Co-Immunoprecipitation

Cell lysis was performed in ELB buffer (160mM NaCl, 5mM EDTA, 50mM HEPES (pH7.5), 0.1% NP-40) for 20 minutes on ice, followed by centrifugation at 13,200 xg for 20 minutes at 4 °C. Protein was quantified using Pierce™ BCA Protein Assay (Thermo Fisher Scientific) according to the manufacturer’s instruction. 600 μg of lysate was diluted in IP buffer (150mM NaCl, 1mM EDTA, 50mM HEPES (pH7.5), 1mM DTT, 2.5mM EGTA, 10% Glycerol) to a total volume of 1 mL and incubated with 25 μL of HA-Trap Magnetic Agarose beads (ChromoTek) under rotation at 4 °C overnight. The beads were then washed 3x in IP buffer and 2x in PBS. Proteins were eluted with 2x SDS loading buffer and heated at 95 °C for 10 minutes, prior to western blot analysis.

### Western Blot

#### Sample Preparation

##### Whole Cell

RIPA buffer combined with a protease and phosphatase inhibitor cocktail was used to lyse the cells for 20 minutes on ice. After incubation, samples were centrifuged at 16,100xg for 10 minutes at 4°C. Supernatants were collected and quantified using the Pierce™ BCA Protein Assay (Thermo Fisher Scientific) according to the manufacturer’s instruction. LDS sample buffer (4X) at 1X final concentration and 2.5% β-Mercaptoethanol were added to 15 µg of protein and heated at 90°C for 10 minutes before Western blot analysis.

##### Subcellular Fractionation

Subcellular fractionation was performed using the Subcellular Protein Fractionation Kit for Cultured Cells (Thermo Fisher Scientific), according to the manufacturer’s instructions. Chromatin fraction was quantified using the Pierce™ BCA Protein Assay (Thermo Fisher Scientific) by following manufacturer’s instructions. LDS sample buffer (4X) at 1X final concentration and 2.5% β-Mercaptoethanol were added to 15 µg of protein and heated at 90°C for 10 minutes before Western blot analysis.

For lambda phosphatase protein (LPP, New England Biolabs) treatment samples were incubated with 2 µL LPP, 1.5 µL Protein MetalloPhosphatases (PMP) and 1.5 µL MnCl_2_ buffers for 1 hour at 30 °C prior to preparation for Western blot analysis.

##### SDS-PAGE

Proteins were separated in a 10% SDS-PAGE pre-cast gel at 150V in 1X MOPS buffer, using PowerEase 90W. Precision Plus Dual Colour ladder was used as a protein size marker.

##### Western Blotting

Separated proteins were transferred from the gel onto a nitrocellulose membrane in a Bio-Rad Criterion™ Blotter tank using wet transfer method. Transfer was performed at 4°C for 1 hours at 100V. Ponceau stain was used to visualise the protein bands after transfer. Membranes were blocked using 5% milk (Serva) in TBST for 1 hour prior to overnight incubation in the appropriate primary antibody at 4°C. Antibodies were diluted in TBST. After incubation and 3 washes of 10 minutes in TBST, membranes were incubated with Horseradish Peroxide (HRP) conjugated secondary antibody for 1 hour at room temperature. Protein bands were visualised on X-ray films using Pierce ECL Chemiluminescent Substrate.

Chromatin Fractionation samples were blotted using LI-COR fluorescent secondary antibodies and visualised on the LI-COR Odyssey classic imager system. Quantification of western blot bands was performed using FIJI software (analyse, gels function).

##### Phos-Tag Western Blotting

ASCL1 phosphorylation post *in vitro* kinase assay was determined using an 8% acrylamide gel polymerised with 20 µM Phos-tag reagent (WAKO) and 40 µM MnCl_2_. After running and before transfer, phos-tag gels were washed three times for 10 min each with transfer buffer (25mM Tris-HCl, 190mM glycine, 20% methanol) supplemented with 10 mM EDTA, followed by a final wash with transfer buffer only. All other steps were followed as described above.

#### RNA sequencing

##### Sample Preparation and Sequencing

For RNA sequencing of IMR^ASCL1^ and BE^ASCL1^ cells were treated with 1 µg/mL doxycycline for 24 hours was performed as described in ^74^. Briefly, RNA was isolated from the following four samples in five biological replicates: IMR^ASCL1^ untreated, IMR^ASCL1^ dox treated, BE^ASCL1^ untreated, BE^ASCL1^ dox treated using RNeasy mini kit (Qiagen) and on-column DNase treatment to remove genomic DNA.

In detail, libraries were prepared from isolated RNA (1 μg) from each sample. RNA was quantified using Qubit BR RNA kit (Invitrogen), and the integrity of RNA was determined using Tape station to be above 9.0 RIN (Agilent). Strand-specific mRNA sequencing libraries were prepared using Illumina Stranded mRNA Library Prep Kit (Illumina), following the manufacturer’s procedure. The fragments were quantified using KAPA quantification kit (Roche), and the library size was determined using Tape station D1000 (Agilent). Libraries were sequenced on the NovaSeq 6000 in a 150 bp paired-end run to a depth of at least 20 million reads per library. Approximately 40-60 million 150-bp paired reads were achieved for each sample.

##### RNA sequencing Bioinformatics Analysis

Reads were trimmed to remove adaptors and to a quality score of 20 with trim galore v0.6.4_dev before aligning to the hg38 genome using STAR v2.6.1d^75^, using quantMode reverse to obtain raw gene counts.

For each cell line, differential expression analysis was conducted using DESeq2 with lfcShrink using apeglm for the shrinkage estimator^38,76^. Genes with counts fewer than the total number of samples (20 counts) were removed before identifying significantly differentially expressed genes (adjusted p value < 0.05 and log_2_FC > 0.5 or log_2_FC < −0.5). Data was vst normalised in DESeq2 for downstream analysis. Significantly changing genes were used for subsequent heatmap plotting. Normalised gene expression (counts per million) in all samples was subject to z-score scaling across both cell lines, prior to visualisation using ComplexHeatmap^77^.

Neuronal or Cell Division gene signatures were generated by selecting related ontology terms from those significantly enriched upon ASCL1 overexpression in either cell line. Ontology terms containing “neur”, “synap”, “axon” were filtered for the Neuronal gene signature, and “mito”, “cycl” for the Cell Division gene signature. Ontology terms containing the word “negative” were excluded from both. List was then manually checked and corrected to only contain relevant GO-terms. The full list for each gene signature is available in the supplementary materials.

#### ASCL1 ChIP sequencing

##### Samples Processing and Sequencing

ASCL1 ChIP sequencing of IMR^ASCL1^ and BE^ASCL1^ cell lines treated with 1 µg/mL doxycycline for 24 hours was performed as described in ^33^. The experiments were performed with four biological replicates for each cell line. Briefly, 20 x 10^6^ cells were fixed with 1% formaldehyde for 10 mins and quenched with 1.5 mM Glycine. The cell pellet was washed twice with 1X PBS supplemented with 1 mM PMSF, harvested by scraping followed by centrifugation and stored at −80⁰C. The cell pellets were lysed using a series of lysis buffers as described in ^78^ to isolate the nuclei, and the DNA was sheared to 100-500 bp fragments using the Bioruptor pico sonicator (Diagenode). 10 µg of ASCL1 antibody (ab74065; Abcam) was used to precipitate bound DNA from around 750 µg of sheared chromatin. Cross-linking was reversed, and ChIP DNA was purified using the Qiagen PCR purification kit (Qiagen) and quantified with the Qubit HS assay kit (Invitrogen). ChIP sample libraries were generated from four biological replicates, and the respective input samples were pooled to generate sequencing library, generated using the NEBNext® Ultra™ II DNA Library Prep Kit (Illumina). The amplified libraries were quantified using the KAPA quantification kit (Roche), and the library size was determined using Tape station D1000 (Agilent). Libraries were pooled and sequenced on the NovaSeq 6000 in a 150 bp paired-end run to a depth of at least 25 million mapped reads per sample.

##### ChIP sequencing Bioinformatics Analysis

Reads were trimmed to remove adaptors and to a quality score of 20 with trim galore v0.6.4_dev before aligning to the hg38 genome using bowtie2 v2.4.1^79^ snakemake wrapper 0.71.1/bio/bowtie2/align. Duplicate reads marked but not removed, and peaks called with macs2 v2.2.7.1, snakemake wrapper 0.66.0/bio/macs2/callpeak -f BAMPE -g hs --SPMR --qvalue 0.05 --keep-dup all.

The DiffBind Bioconductor package^80^ was used to create a consensus set of peaks for each cell line, considering peaks present in at least three out of four replicates. These were combined into an overall consensus peak set, which was used for differential binding analysis in DiffBind. Prior to generating the consensus lists, a blacklist (DBA_BLACKLIST_HG38) and a calculated greylist, generated using input samples with the dba.blacklist command, were applied to the data. Samples were normalized according to library size. Significantly differential ASCL1 peaks between the cell lines were identified using DESeq2 within DiffBind, with significance defined by an adjusted FDR <0.01 and log2FC > |1|. Common (non-significant) group was defined by an FDR > 0.5.

Peak annotation was performed using the ChIPseeker package^81^, with nearest gene annotation limited to a 50 kb distance.

To visualise genomic loci of ASCL1 peaks, tag density and average profile plots were generated using deeptools computeMatrix, plotHeatmap and plotProfile, on RPKM normalised (deeptools^82^ bamCoverage) and averaged (WiggleTools^83^) bigwig files.

Motif analysis was performed using the HOMER findMotifsGenome.pl command^84^ with the - size parameter set to 100. For direct comparisons between two sets of sequences, one set was used as the background in the findMotifsGenome.pl command.

##### Gene Ontology Analysis

Gene ontology analysis was performed using compareCluster from clusterProfiler package^85^ in R. The following parameters were selected OrgD-b=”org.Hs.eg.db”, pAdjustMethod=”fdr”, minGSSize=20, maxGSSize=500, ont=”BP”, pvalueCutoff=0.01.

##### Quantification and statistical analysis

Statistical analyses were performed using Prism software as stated in the figure legends (ns, non-significant, ∗, P ≤ 0.05; ∗∗, P ≤ 0.01, ∗∗∗, P ≤ 0.001). The standard deviation was calculated from at least three independent experiments. n numbers are stated in the figure legends. Statistical analysis of differentially expressed genes from RNA sequencing was performed using DESeq2, and of differential ASCL1 peaks from ChIP sequencing using DESeq2 with DiffBind in R v.4.3.1. Whole proteome and ASCL1 qPLEX-RIME data was statistically analysed using limma^86^ within the qPLEXanalyzer Bioconductor package^44,73^. Details of these analyses are described in their respective methods sections. Biological replicates were considered as different passage of the same cell line seeded in independent experiments.

## Acknowledgments

We would like to thank Kirsty Ferguson, Roberta Azzarelli, Frances Connor, William Beckman, and all Philpott lab members for helpful discussions. This research was supported by the CRUK Cambridge Institute Proteomics Core Facility and Cambridge NIHR BRC Cell Phenotyping Hub. We would also like to thank the Cambridge Stem Cell Institute Imaging Core Facility, and the Research Instrumentation and Cell Services facility and the Bioinformatics facility at the CRUK Cambridge Institute. We would like to thank Ahmed Abou Tayoun and Maha ElNaofal (Al Jalila Genomics Center of Excellence, Al Jalila Children’s Specialty Hospital, Dubai, United Arab Emirates) for sequencing support. Thank you to Prof. Deborah Tweedle for generously supplying neuroblastoma cell lines.

This work was supported by the Cancer Research UK grant A25636 (to A.P., L.M.W., D.M., R.D.), Wellcome Trust Investigator award 212253/Z/18/Z (to A.P., L.M.) and by Neuroblastoma UK (to D.M. and A.P). L.M. acknowledges support from Cancer Research UK MRes/PhD Studentship Award and J.L.B. from Wellcome Trust MRes/PhD Studentship Award. F.R.A. acknowledges support from Mohammed Bin Rashid University of Medicine and Health Sciences (MBRU-CM-RG2023–08), Al Jalila Foundation (to R.L.G, R.R. and F.R.A.), The Terry Fox Foundation, and MBRU ALMAHMEED Collaborative Research Award (ALM1909). J.S.C. acknowledges support from the University of Cambridge, Cancer Research UK core funding (grants A20411, A31344, A29580, and DRCPGM\100088) and Hutchison Whampoa. Core support was provided from the Wellcome Trust 203151/Z/16/Z and the UKRI Medical Research Council MC_PC_17230 (to L.M., L.M.W., J.L.B., D.M., R.D. and A.P.). For the purpose of open access, the author has applied a Creative Commons Attribution (CC BY) licence to any Author Accepted Manuscript version arising from this submission.

## Supplemental information

Table S1. Excel file listing the gene ontology terms and genes making up the neuronal gene signature

Table S2. Excel file listing the gene ontology terms and genes making up the cell division gene signature

Table S3. Excel file of the ASCL1 interactors in dox treated BE^ASCL1^ versus IMR^ASCL1^ identified with ASCL1 qPLEX-RIME.

Table S4. Excel file of the total proteome analysis of dox treated BE^ASCL1^ versus IMR^ASCL1^.

## Inclusion and diversity

We support inclusive, diverse, and equitable conduct of research.

## Author contributions

Conceptualization, L.M. and A.P.; Methodology, L.M., L.M.W., A.P.; Guidance on methodology, E.K.P. and J.S.C.; Formal Analysis, L.M.; Investigation: L.M., E.K.P., R.L.G., R.R., L.M.W., J.L.B., D.M., R.D.; NGS data processing, L.M.W.; bioinformatic analysis, L.M.; Visualisation, L.M.; writing – original draft, reviewing & editing, L.M., and A.P.; funding acquisition, F.R.A. and A.P.; project administration, A.P.; supervision, F.R.A., J.S.C. and A.P.

## Key resources table

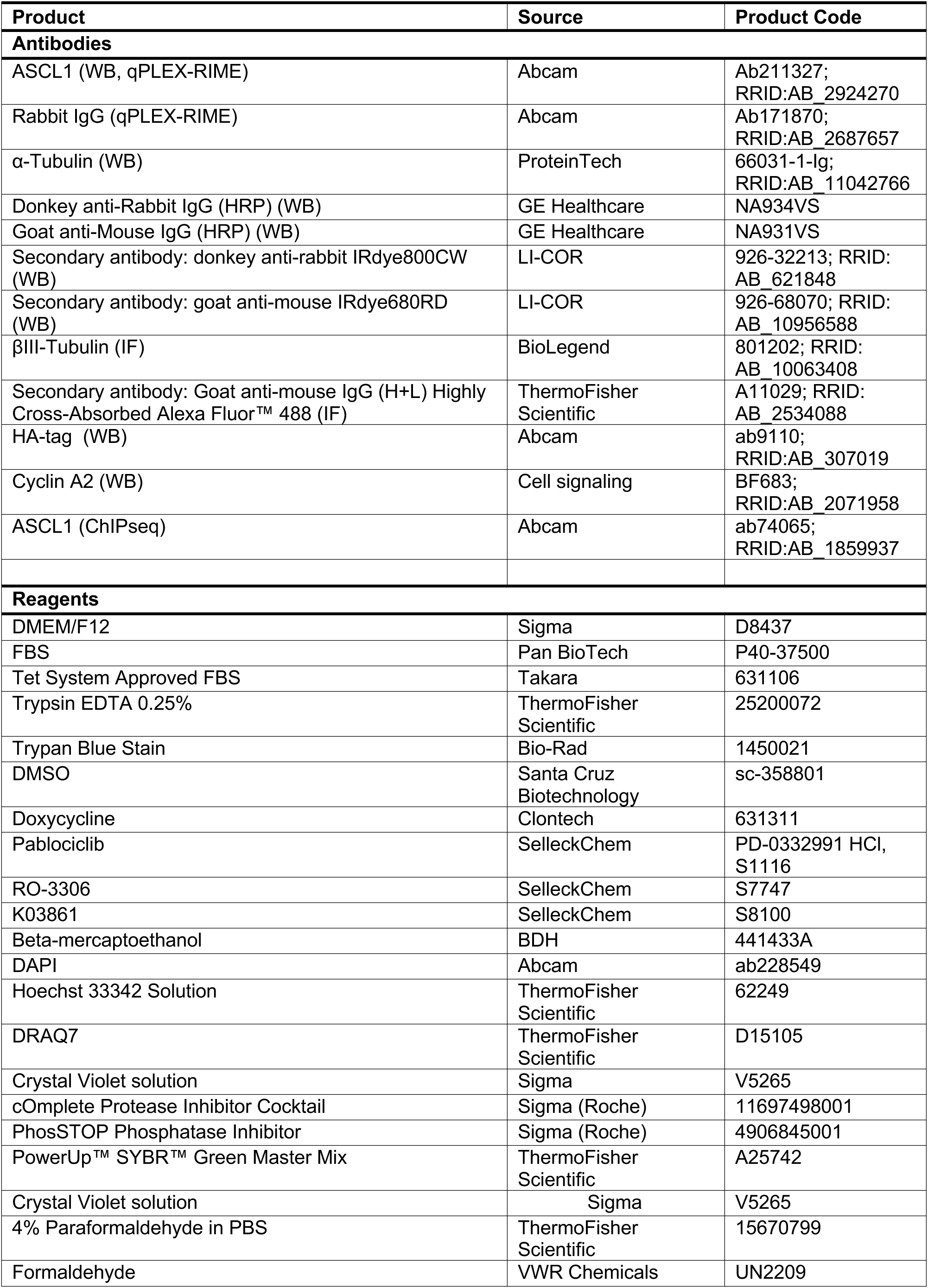
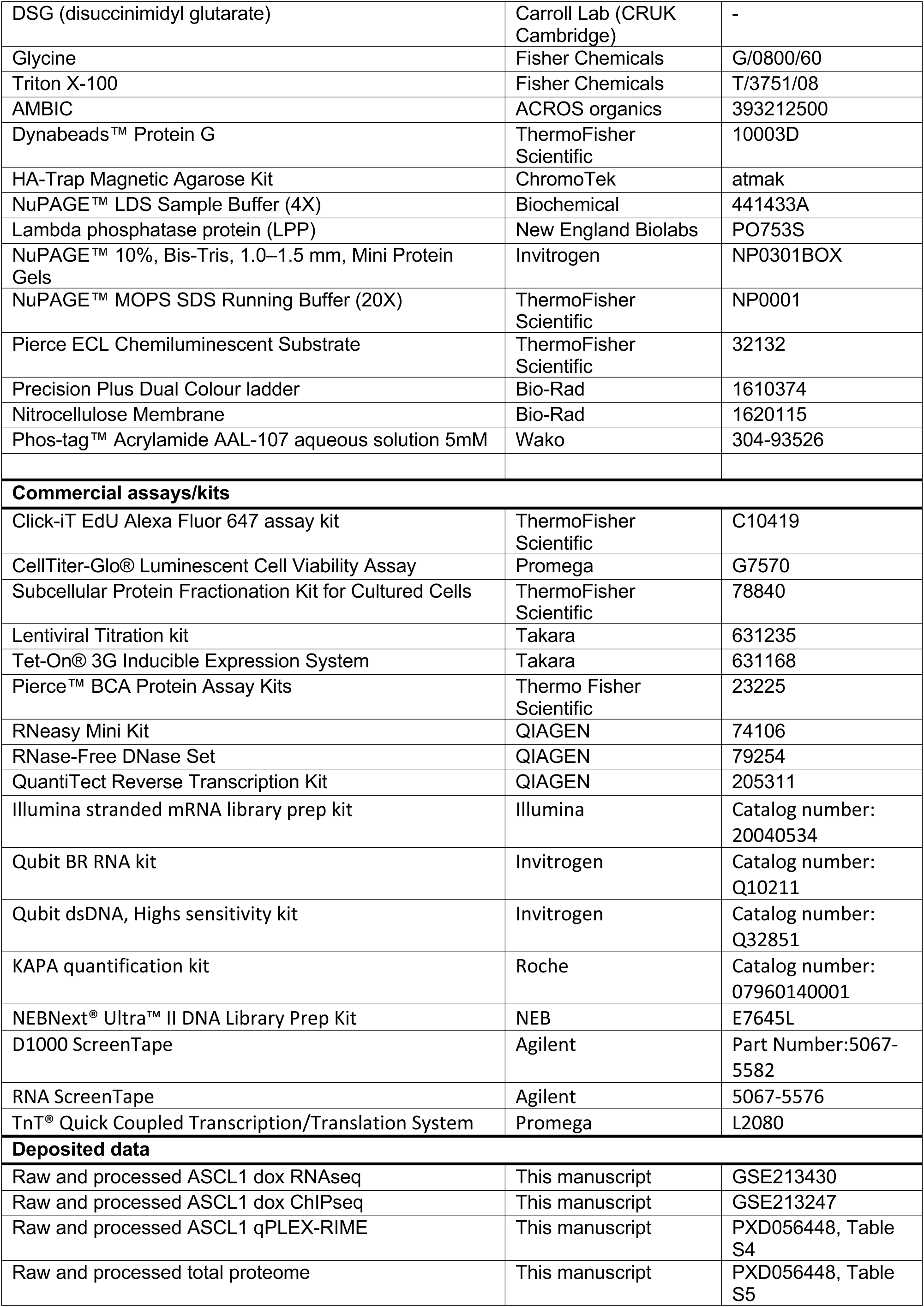
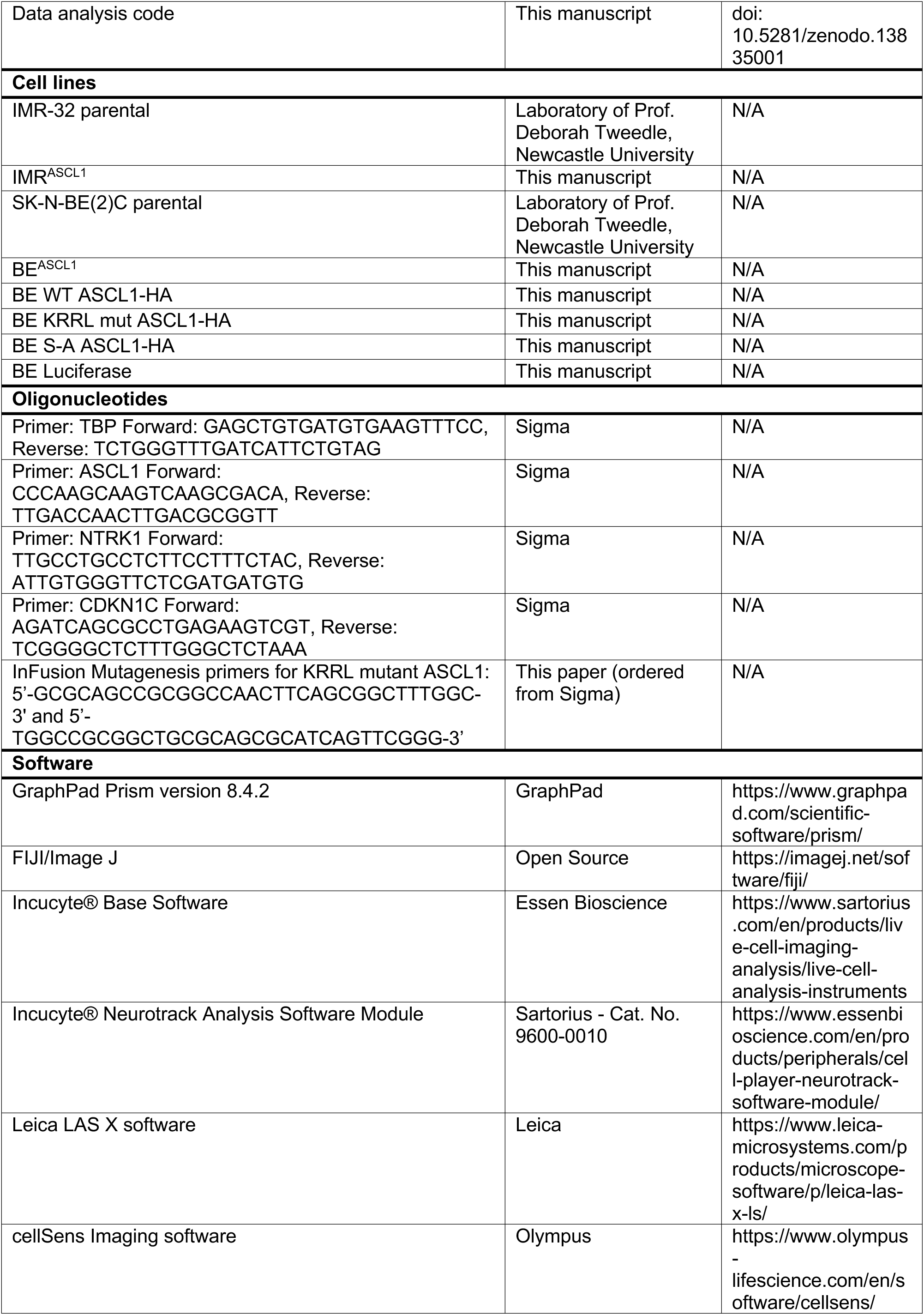
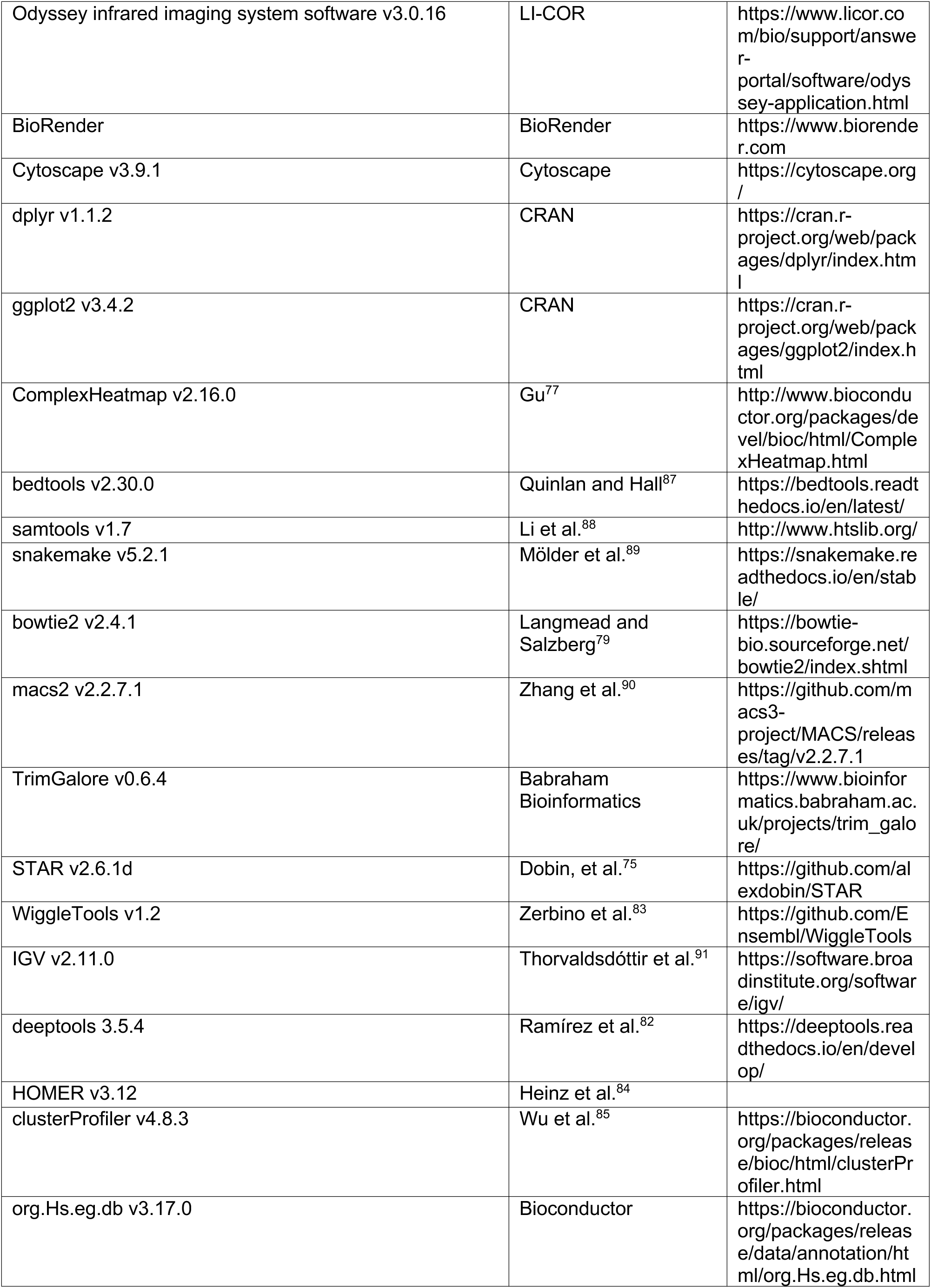
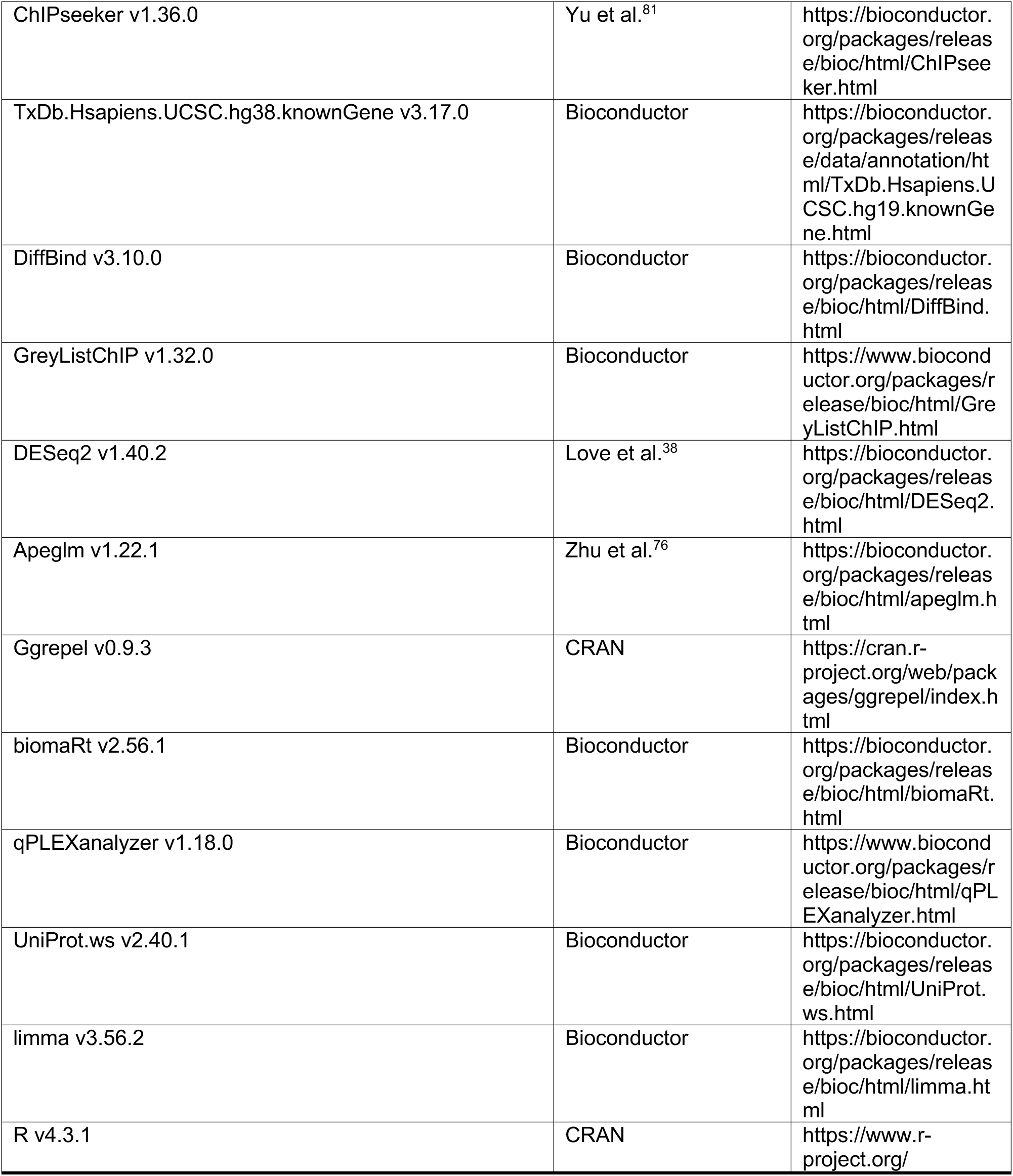

## Notes

**Conflict of interest statement:** The authors declare no potential conflicts of interest.

### Competing Interest Statement

The authors have declared no competing interest.

